# Multi-cellular communities are perturbed in the aging human brain and Alzheimer’s disease

**DOI:** 10.1101/2020.12.22.424084

**Authors:** Anael Cain, Mariko Taga, Cristin McCabe, Gilad Green, Idan Hekselman, Charles C. White, Dylan I. Lee, Pallavi Gaur, Orit Rozenblatt-Rosen, Feng Zhang, Esti Yeger-Lotem, David A. Bennett, Hyun-Sik Yang, Aviv Regev, Vilas Menon, Naomi Habib, Philip L. De Jager

**Author notes:** Current address: Genentech, 1 DNA Way, South San Francisco, CA. Equal contribution (Anael Cain, Mariko Taga). Equal contribution (Aviv Regev, Vilas Menon, Naomi Habib, Philip L. De Jager). Corresponding Authors: Philip L. De Jager, Naomi Habib, Vilas Menon.

## Abstract

The role of different cell types and their interactions in Alzheimer’s disease (AD) is an open question. Here we pursued it by assembling a high-resolution cellular map of the aging frontal cortex by single nucleus RNA-seq of 24 individuals with different clinicopathologic characteristics. We used the map to infer the neocortical cellular architecture of 638 individuals profiled by bulk RNA-seq, providing the sample size necessary for identifying statistically robust associations. We uncovered diverse cell populations associated with AD, including inhibitory neuronal subtypes and oligodendroglial states. We further recovered a network of multicellular communities, each composed of coordinated subpopulations of neuronal, glial and endothelial cells, and found that two of these communities are altered in AD. Finally, we used mediation analyses to prioritize cellular changes that might contribute to cognitive decline. Thus, our deconstruction of the aging neocortex provides a roadmap for evaluating the cellular microenvironments underlying AD and dementia.

## INTRODUCTION

Over the past decade, our understanding of the molecular landscape of Alzheimer’s disease (AD) has advanced rapidly as new experimental and analytic methods synergized to uncover pathways that contribute to the sequence of events that lead to AD dementia. While genetic studies have created a strong foundation of selected loci and genes implicated as causal in susceptibility to AD and its endophenotypes, transcriptomic analyses have sketched a broader landscape of molecular changes that capture disease dynamics and the state of the target organ. However, most prior efforts profiled RNA at the bulk tissue level (*e.g.* ^1^), averaging expression measures across a myriad of cell types and states, which obscured finer distinctions and contributions from cell subtypes found at low frequency in the brain.

Recent studies that profiled single nuclei from post-mortem brain tissue of healthy and AD individuals have uncovered specific cell types with different signatures and proportions in AD, especially in certain subsets of microglia, oligodendrocytes and astrocytes^2–6^. While such studies hint at substantial inter-individual variability, their limited sample size and moderate number of profiled nuclei per subject yields an incomplete picture of the architecture of the aging neocortex and limits the statistical power of association analyses. In addition, while most studies focus on a cell-intrinsic view, cellular interactions and dependencies between different cell types and subtypes in the aging and AD brain remain under-explored but could play crucial role in disease pathology.

Several key questions remain open: (1) How distinct is the cellular architecture in AD brains? (2) Are changes in certain cellular subsets coordinated with or independent of one another? (3) How do changes in the cellular architecture relate to the causal chain of events leading to AD? Addressing these questions currently requires several new tools including methods to relate detailed maps of single nucleus profiles from select individuals to large cohorts of deeply phenotyped individuals with sufficient size to complete statistically robust association studies, as well as approaches that characterize both individual cells and multi-cellular communities.

Here, we have deployed a combined approach that integrates single nucleus RNA-seq profiling (snRNA-seq) of the dorsolateral prefrontal cortex (DLPFC) tissue from a structured subgroup of 24 well-characterized human subjects, together with bulk RNA profiles of a statistically well-powered set of 638 deceased subjects^1^. Each of these subjects was a participant in either the ROS (Religious Order Study) or the MAP (Memory and Aging project): two harmonized longitudinal studies of cognitive aging with prospective autopsy and deep cognitive and neuropathologic characterization that are managed by a single group of investigators at Rush University and are designed to be analyzed together^7–9^. The 24 participants profiled by snRNA-seq span four archetypal categories of older women and men, to ensure the capture of the cellular diversity of this brain region across a range of clinicopathologic states. By contrast, the 638 participants with previously collected bulk RNA-seq profiles^1^ are a relatively random sample of the cohorts and reflect the general distribution of characteristics seen in the older population, enabling robust statistical modeling (**Fig. 1a**). Compared to previous studies^2–4^, our careful selection of donors, deeper sampling of single nuclei profiles, and high library quality yielded an enhanced map of cell subtypes and cell states of the aging cortex. We extended this map with a new computational approach, CelMod, to hierarchically estimate broad cell class and subtype proportions across the 638 bulk RNA-seq profiles. The inferred proportions in a large cohort provided statistical power to identify cell subpopulations associated with pathophysiology of AD. We uncovered an association between cognitive decline and the proportion of specific subpopulations of glial cells, oligodendrocytes, microglia and astrocytes, and subtypes of inhibitory neurons, especially, a relative decrease in somatostatin (SST) neurons. We further used our data to infer a map of multi-cellular communities whose proportions are correlated across individuals, which may reflect micro-environments in the aging brain. Two anti-correlated communities, each displayed opposite associations with cognitive decline in AD and with tau pathology load. Finally, causal modeling suggested that the differences in cell subtypes and signatures occur downstream of the accumulation of tau pathology and might mediate downstream effects that accelerate cognitive decline. Our model can inform further investigation and therapeutic development, by identifying those cellular factors that may most proximally and directly contribute to loss of cognitive function with advancing age and AD.

**Figure 1.**
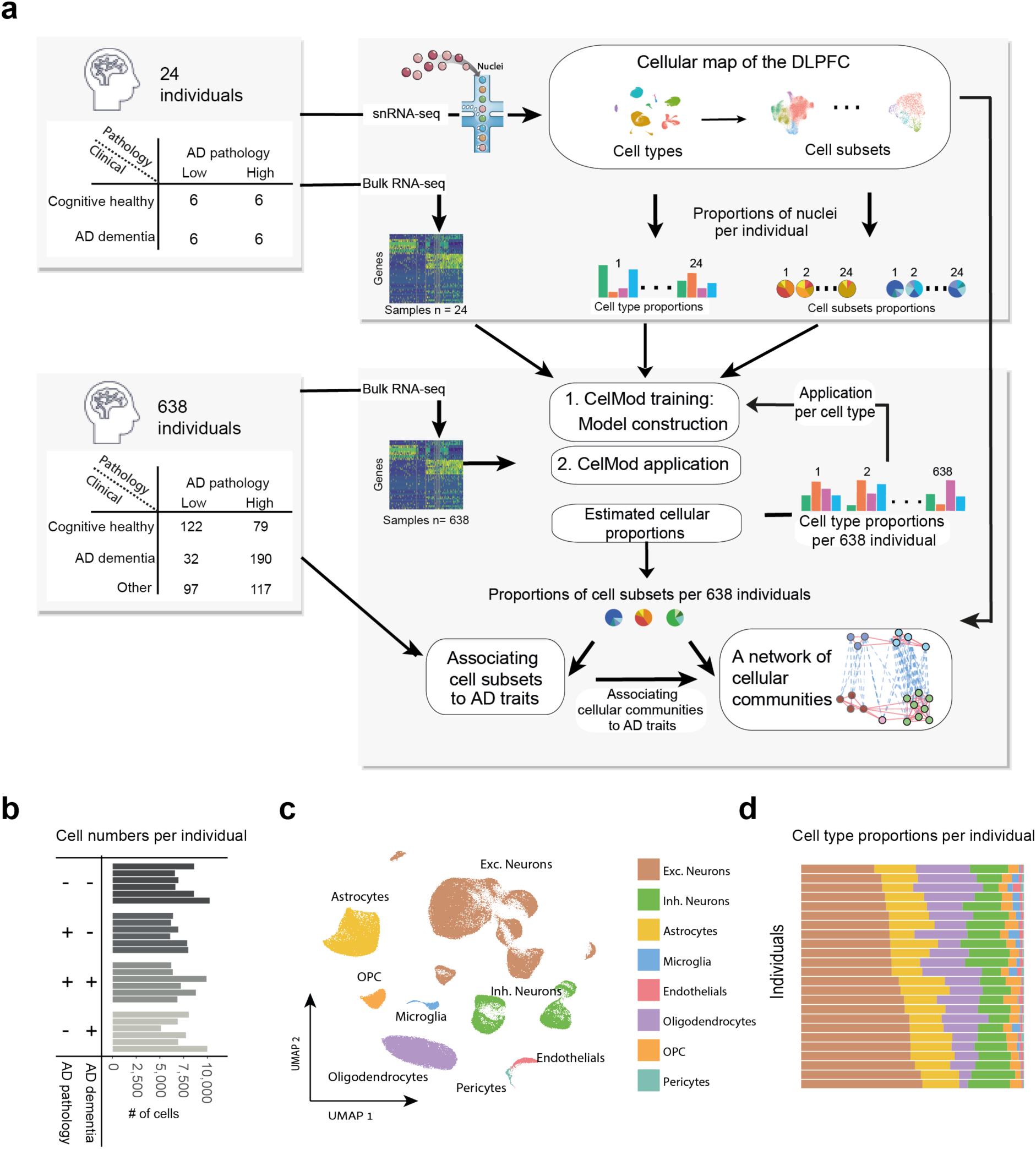
A cellular-molecular map of the human aging DLPFC in 24 cognitively healthy and Alzheimer’s Disease (AD) individuals. (**a**) Overview of the experimental scheme and analysis. 24 individuals with clinicopathologic characteristics were profiled by single nucleus RNA-seq (snRNA-seq) to generate a high resolution *cellular map* of the aging DLPFC brain region, and used as input to our CelMod deconvolution algorithm to iteratively estimate *cellular compositions* in 638 individuals. Network analysis uncovered *cellular communities*, cell subsets coordinately varying across individuals, and statistical modeling *associated AD traits* to cell subsets and to cellular communities. (**b**) High quality 172,659 nuclei libraries generated across 24 post-mortem samples of the DLPFC brain region of aging individuals. The number of cell profiles for each individual, ordered by the four major archetypes of the aging population: reference (non-impaired individuals with minimal AD pathology), resilient (cognitively non-impaired with a pathologic diagnosis of AD), AD group (both clinical and pathologic AD), and clinical-AD-only (AD dementia with minimal AD pathology). (**c**) Umap embedding of 172,659 single-nuclei RNA profiles from the DLPFC brain region of 24 individuals; colored by cell type. (**d**) Diversity of cell type proportions across individuals. The proportion of cell types, color coded as in (c), for each individual (rows).

## RESULTS

### Inter-individual diversity drives cellular landscapes in the aging DLPFC

To build a map of the aging DLPFC (BA9), we generated snRNA-seq profiles from frozen tissue samples obtained from 24 ROSMAP^8,9^ participants with an average age of 87.9 years at the time of death; demographic details are presented in **Supplementary Table 1**. To sample a wide variety of cell subtypes and states, we selected participants that represent four major archetypes of the aging population (**Fig. 1a, Supplementary Table 1**): (1) a reference group of cognitively non-impaired individuals with minimal AD pathology, (2) a resilient group of cognitively non-impaired individuals with a pathologic diagnosis of AD, (3) an AD group who fulfill diagnostic criteria for both clinical AD dementia and pathologic AD, and (4) a clinical-AD-only group diagnosed with AD dementia but only minimal AD pathology post-mortem. Each group consists of 6 individuals (50% men/women). This sampling strategy ensured that we capture a wide range of cellular signatures across the different pathological and clinical manifestations of the disease within the 24 individuals profiled by snRNA-seq. We retained 172,659 DLPFC nuclei profiles for analysis, with an average of 7,194 nuclei per participant after doublet filtering, background subtraction, and quality control. (**Fig. 1b, Extended Data Fig. 1a, Methods**).

Unsupervised clustering identified distinct groups of nuclei spanning all 8 major expected cell types (**Fig. 1c, Extended Data Fig. 1b,c, Supplementary Table 2**). Nuclei from each group were present in each individual, but with considerable inter-individual variation in their relative proportions (**Fig. 1d**), which led us to explore the inter-individual diversity within each cell type across subtypes and cellular states.

### A high-resolution cell atlas of the aging neocortex

We analyzed profiles from each of the 8 major cell types separately (**Fig. 2, Fig. 3, Extended Data Fig. 2-3**), to explore finer distinctions in cellular diversity, including cell subtypes, states and expression programs in the neocortex. Overall, our single nucleus-derived cell subtype model is consistent with earlier, lower-resolution models of microglia, astrocytes, oligodendrocytes and endothelial cell subtypes, as well as with previous neuronal clustering efforts (**Extended Data Fig. 4**) showing the consistency of data generated independently from frozen cortex^2,3,5,10^. Thus, the overall population structure of cells is well captured in our data and allows us to evaluate the interrelation of each cell subset with other cell types and states.

**Figure 2.**
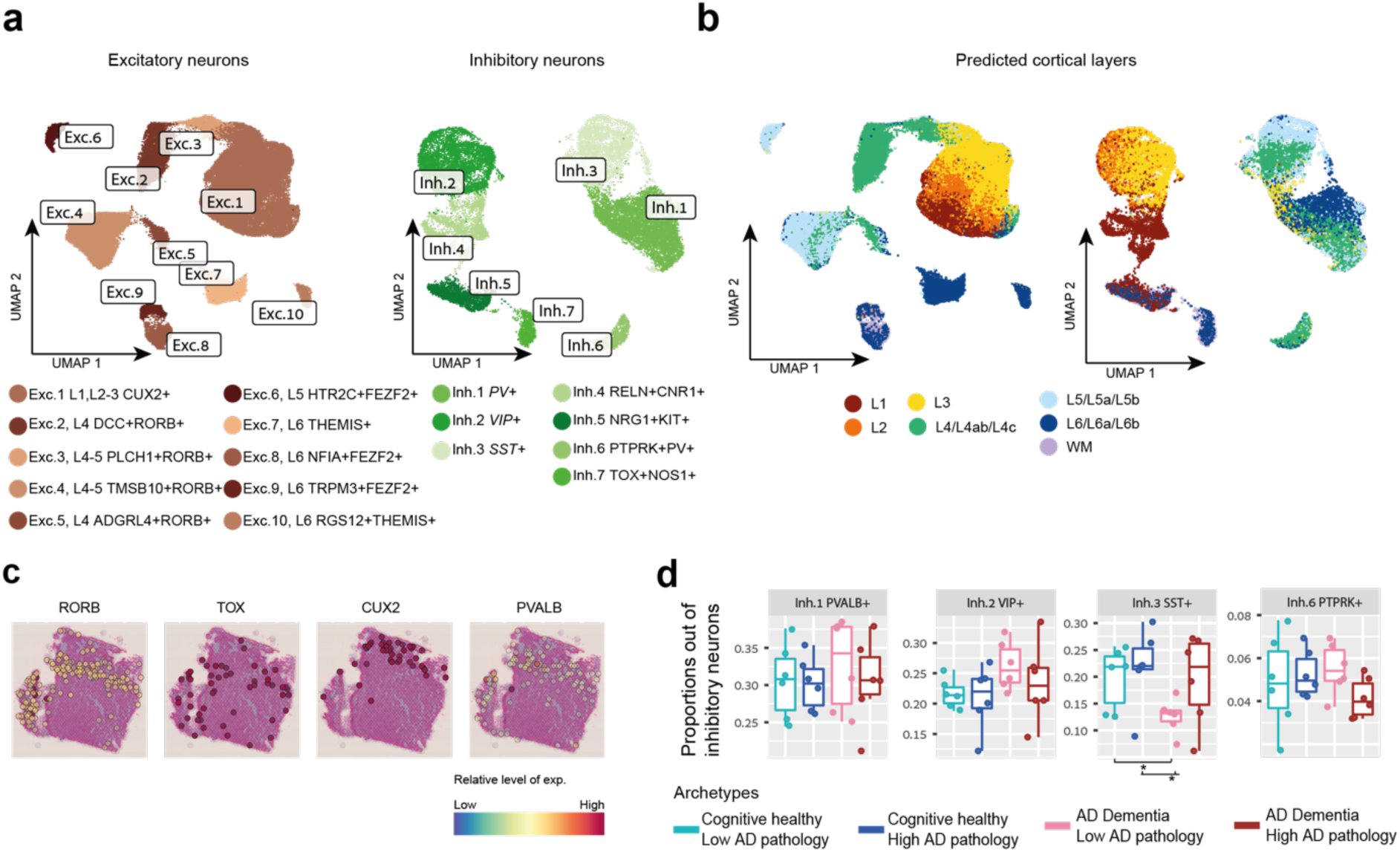
Diversity of neuronal subtypes across layers in the aging DLPFC. **(a)** Neuronal diversity in the DLPFC. Left: Umap embedding of excitatory (74,999 cells, 10 clusters) and Right: inhibitory neuronal subtypes (24,938 cells, 7 clusters), colored by clusters capturing distinct subtypes. Annotated by known neuronal markers (**Extended Data Fig. 2a**) and mapped to previous annotations^11^. (**b**) Neuronal diversity in the DLPFC captures neurons at the various cortical layers. Umaps of neuronal subtypes colored by predicted cortical layers according to a classifier applied to annotated RNA profiles from Allen Brain Atlas^11^ (**Methods**) (**c**) Visium spatial transcriptomics highlights a layer pattern in DLPFC slices using cortical neuronal markers (RORB, CUX2, TOX, PVALB). (**d**) Proportion of SST GABAergic neuronal subtype varies in association with cognitive decline. Box plots of the proportion of three GABAergic subtypes (out of total GABAergic neurons) across the four major archetypes of the aging population: reference group, pathological AD only (resilient), clinical and pathological AD, and clinical AD only. Box, 25% and 75% quantiles; line, median; Dots, individual samples.

**Figure 3.**
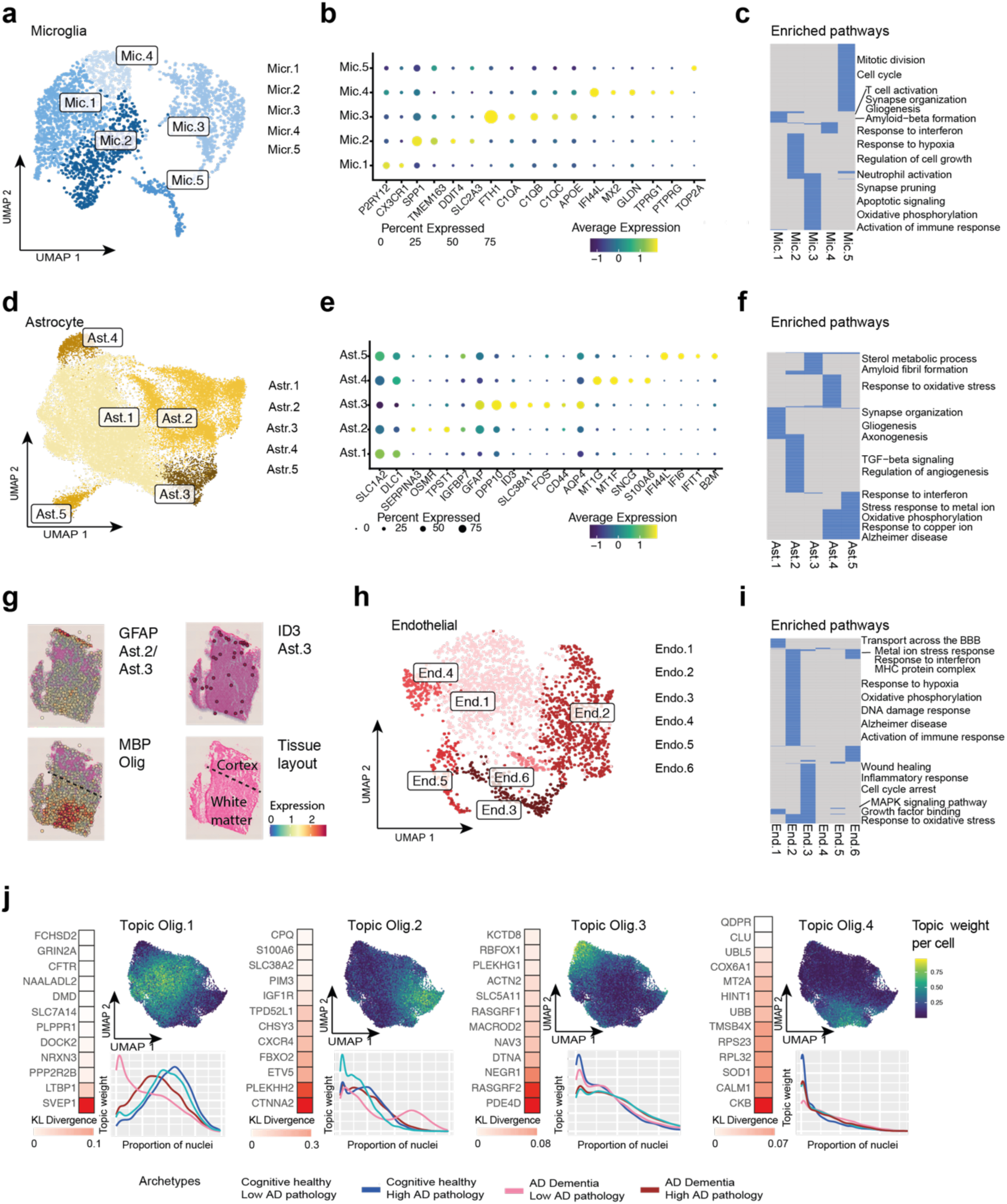
Diverse cell states of glial and endothelial cells in the aging DLPFC. **(a, d, h)** Diversity of non-neuronal cell states in the aging DLPFC. Umap embedding of (a) microglia 2,837 cells, (d) astrocytes with 29,486cells and (h) endothelial with 2,296 cells, colored by clusters capturing distinct cell states (**b, e**) Dot plot of the mean expression level in expressing cells (color) and percent of expressing cells (circle size) of selected marker genes (columns) across microglia (b) or astrocytes (e) subsets (rows). (**c, f, i**) Enriched pathways (hypergeometric test, FDR < 0.05, blue color) in up-regulated gene signatures for each subset of microglia (c), astrocyte (f), or endothelial (i) cells (columns). (**g**) Visium spatial transcriptomics showing position of Ast.3 cells by marker gene ID3 and reactive astrocytes marker GFAP. MBP oligodendrocyte marker marks white matter border. Bottom right: schematics of the tissue. Additional tissues and genes at **Extended Data Fig. 5**. (**j**) Continuum of expression programs in oligodendrocyte cells inferred by topic modeling ^56^. For each topic model: Umap embedding of oligodendrocytes cells, colored by the weight of each topic per cell (right); The top scoring genes (colored by the score), computed as the Kullberg-Leibler (KL) divergence between the expression level and the topic’s weight across cells (red color scale, left); and The cumulative distribution function of topic weights for cells split by the sample of origin to four major archetypes of the aging population (as in **Fig.2 d**).

Among neurons, we identified 10 excitatory and 7 inhibitory subsets (74,999 and 24,938 nuclei respectively, **Fig. 2a, Extended Data Fig. 2a-c**), capturing the diversity of the neocortex. Each subset expressed unique marker genes, including known markers, such as the inhibitory subtype markers somatostatin (SST, in Inh.3 cluster) or parvalbumin (PV, in Inh.1 cluster) (**Extended Data Fig. 2a**). Using a machine learning classifier (Methods), we mapped our subsets to previously annotated neuronal subtypes^11^, showing that our clusters aligned well with previous annotations and captured transcriptionally and spatially distinct pyramidal neurons across the different cortical layers as well as inhibitory neuronal subtypes (**Fig. 2a**). Distinct cell clusters were associated with different cortical layers (**Fig. 2b**), which we confirmed using spatial transcriptomics (**Fig. 2c, Extended Data Fig. 2d**). Notably, the proportions of the SST (Inh.3) neuronal cells was nominally lower in individuals exhibiting cognitive decline (**Fig. 2d**), although the small sample size of this dataset was underpowered for meaningful statistical analysis.

Similarly, non-neuronal cell types including astrocytes, microglia and endothelial cells (**Methods, Fig. 3, Extended Data Fig. 3, Supplementary Table 2**), each partitioned to subsets (clusters) that were robust to regression of technical parameters (RIN and batch) and to sex differences (**Methods, Extended Data Fig. 3b-d**). These subsets aligned with previous annotations (**Extended Data Fig. 4**). On the other hand, clustering analysis did not identify robust subdivisions among oligodendrocyte progenitor cells (OPCs) and pericytes (possibly due to their relatively lower numbers); oligodendrocytes also did not have clearly discrete subgroupings, leading us to use a different approach to model this abundant cell type: we assessed co-varying gene programs (topic models, below).

The 2,837 microglial nuclei were partitioned into 5 major subsets (**Fig. 3a**). Microglia subsets matched expression of known marker genes (**Fig. 3b, Supplementary Table 3**) and distinct pathways (**Fig. 3c**) and mapped clearly to previously defined subsets of scRNA-seq of live human microglia from the DLPFC^6^ (**Extended Data Fig. 4a**). These were annotated as surveilling (Micr.1, expressing P2RY12 and CX3CR1), stress response/anti-inflammatory (Micr.2, expressing SPP1 and TMEM163), enhanced redox (Micr.3, expressing C1Q and APOE), interferon-response (Micr.4, expressing IFI44L) and proliferative (Micr.5, expressing TOP2A) microglia. Micr.2 captured two different reacting states seen in living microglia isolated from fresh autopsy tissue of the DLPFC^6^ (**Extended Data Fig. 4a**), probably due to the smaller number of nuclei (compared to cells) available for analysis. However, our snRNA-seq data identified all of the RNA signatures present in live sorted microglia, suggesting that neither isolation protocol misses specific microglial states or subtypes found in the other approach (we find the same subtypes whether we use a nuclear or live cell approach). Furthermore, while there are differences in individual gene levels in RNA profiles from microglial nuclei *vs*. cells (consistent with earlier reports^12,13^), the nuclear profiles show expression of known markers of microglial subtypes and microglial activation (e.g. SPP1, APOE, TMEM163, **Fig. 3b**), and differences in levels of these genes help to mark some of our microglial clusters from each other, as in the live microglial scRNA-seq dataset (**Supplementary Table 3**).

The astrocytes (29,486 nuclei) included five major subsets, matching known marker genes (**Fig. 3d,e, Supplementary Table 3**), and annotated as: homeostatic protoplasmic-like (Astr.1, highly expressing SLC1A2), non-homeostatic (Astr.2, expressing reactive markers genes GFAP, SERPINA3N and OSMR^14^), meningeal/fibrous-like astrocytes (Astr.3, expressing reactive marker GFAP, ID3^14^ as well as the fibrous/meningeal marker CD44^15^ and enriched in endfeet marker AQP4^16^), Ast.4 (expressing S100A6, MT1A) enriched for oxidative phosphorylation, oxidative and metal ion stress response and Alzheimer’s disease (qvalue <0.05, hypergeometric test, **Fig. 3f**), and a small subset (detected in 2 of the 24 individuals) expressing interferon-response genes (Astr.5, *e.g.* IFI44L, IFI6) (**Fig. 3e,f**). Interestingly, while both Astr.2 and Ast.3 expressed the reactive marker GFAP (**Fig. 3e**), Ast.2 nuclei uniquely expressed genes enriched for axonogenesis, regulation of angiogenesis, and TGF-β signaling, whereas Astr.3 cells were uniquely enriched for sterol metabolic process and amyloid fibril formation genes (qvalue<0.05, **Fig. 3f**). We validated the distinct spatial position of Astr.3 in proximity to the meninges in the neocortex by spatial transcriptomics based on the coexpression of ID3 and GFAP markers (**Fig. 3g, Extended Data Fig. 5**), suggesting these might be interlaminar astrocytes. The Ast.4 profiles expressed markers (e.g. COL5A3, PDE4DIP) previously proposed to have higher expression levels in AD in the human cortex, while the Ast.2 subcluster expressed genes (e.g. SERPINA3) previously proposed to have reduced expression in AD in the human cortex^5^ (**Extended Data Fig. 4b**).

Endothelial cells (2,296 nuclei) included six major subsets, matching recently defined marker genes of the endothelial compartments^17^ (**Fig. 3h, Extended Data Fig. 3e, Supplementary Table 3**): most were from capillary cells (End.1, End.2 and End.3), and the rest annotated as venous (End.4, expressing TSHZ2), arterial (End.5, expressing ARL15) and smooth muscle (End.6, expressing DEPP1) cells. Each of the two major subsets of endothelial capillary cells (End.1 and End.2) expressed genes associated with distinct pathways (qvalue<0.05, hypergeometric test, **Fig. 3i**): including transport across blood−brain barrier and growth factor binding for End.1, and interferon signaling response, oxidative and metal ion stress response, response to DNA damage and Alzheimer’s disease for End.2 (qvalue<0.05, **Fig. 3i**).

### Distinct expression programs underlaying the diversity of aging oligodendrocyte cells

As oligodendrocytes (29,543 nuclei) spanned several gradients of expression, without discreet boundaries, we applied topic modeling using Latent Dirichlet Allocation (LDA)^18–21^ (**Fig. 3j, Supplementary Table 2, Methods**), to recover gene programs (called “topics”) based on co-variation patterns of gene expression across cells. Each gene has a score for each topic, indicating its contribution to the topic, and each nucleus profile has a weight for each topic, indicating the extent to which that topic (program) is present in a particular nucleus. Topics are weighted differently across each nucleus, modeling every cell as a mixture of gene programs. We annotated each topic by their highly scoring genes per topic, using a score based on the Kullberg-Liber (KL) divergence to compare the distribution of topic weights and the expression level of genes across cells^20^ (**Methods, Supplementary Table 3**).

We found four major topics in oligodendrocyte cells (**Fig. 3j, Supplementary Table 2-3**) and their gene assignments, some with potential associations to AD. For example, we found an AD associated cellular adhesion protein (SVEP1^22^) that was highly weighted in topic Olig.1 while the third strongest susceptibility locus for late-onset AD - clusterin (CLU) ^23^ - was highly weighted in Olig.4.

Further, Olig.4 and Olig.2 topics included genes (e.g. *QDPR*) previously reported to have higher expression in cortical oligodendrocytes in AD, while Olig.1 and Olig.3 included genes (e.g. *MOG*) previously proposed to have lower expression in AD in human cortex^5^ (**Extended Data Fig. 4b**

). Moreover, examining the relative distribution of cells from the 24 participants across the four archetype groups suggests that oligodendrocyte signatures are distributed differently among groups, in a manner that is partially mirrored by the distribution of topics (**Fig. 3j**). Thus, such shifts in cellular states may be related to differences in gene programs associated with pathology and cognitive decline; however, the small sample size of these single nucleus data hinders meaningful statistical analysis. We thus subsequently deployed this refined atlas of cell subsets and topics to interrogate a dataset of appropriate size to enable association analyses.

### Deconvolution of cell subtype composition in bulk RNA-seq profiles of 638 individuals

Given substantial heterogeneity in the clinicopathologic characteristics of aging captured by the ROSMAP participants, our 24 “archetypal” participants profiled by snRNA-seq cannot capture the continuous distribution of traits in aging brains of the full ROSMAP studies nor is their number sufficient for robust statistical associations. To overcome the limited statistical power of our sample size, we used our snRNA-seq census of the cellular population structure of the aging cortex to infer cell type proportions in bulk DLPFC RNA profiles from 638 ROSMAP participants (**Supplementary Table 1**). To this end, we leveraged the matching snRNA-Seq and bulk RNA-seq from the DLPFC, available for each of the 24 participants, and **CelMod** (Cellular Landscapes Modeling by Deconvolution), a new method we developed here that builds and validates a model of cell subtype and state proportions from such matched data (**Fig. 4a**).

**Figure 4.**
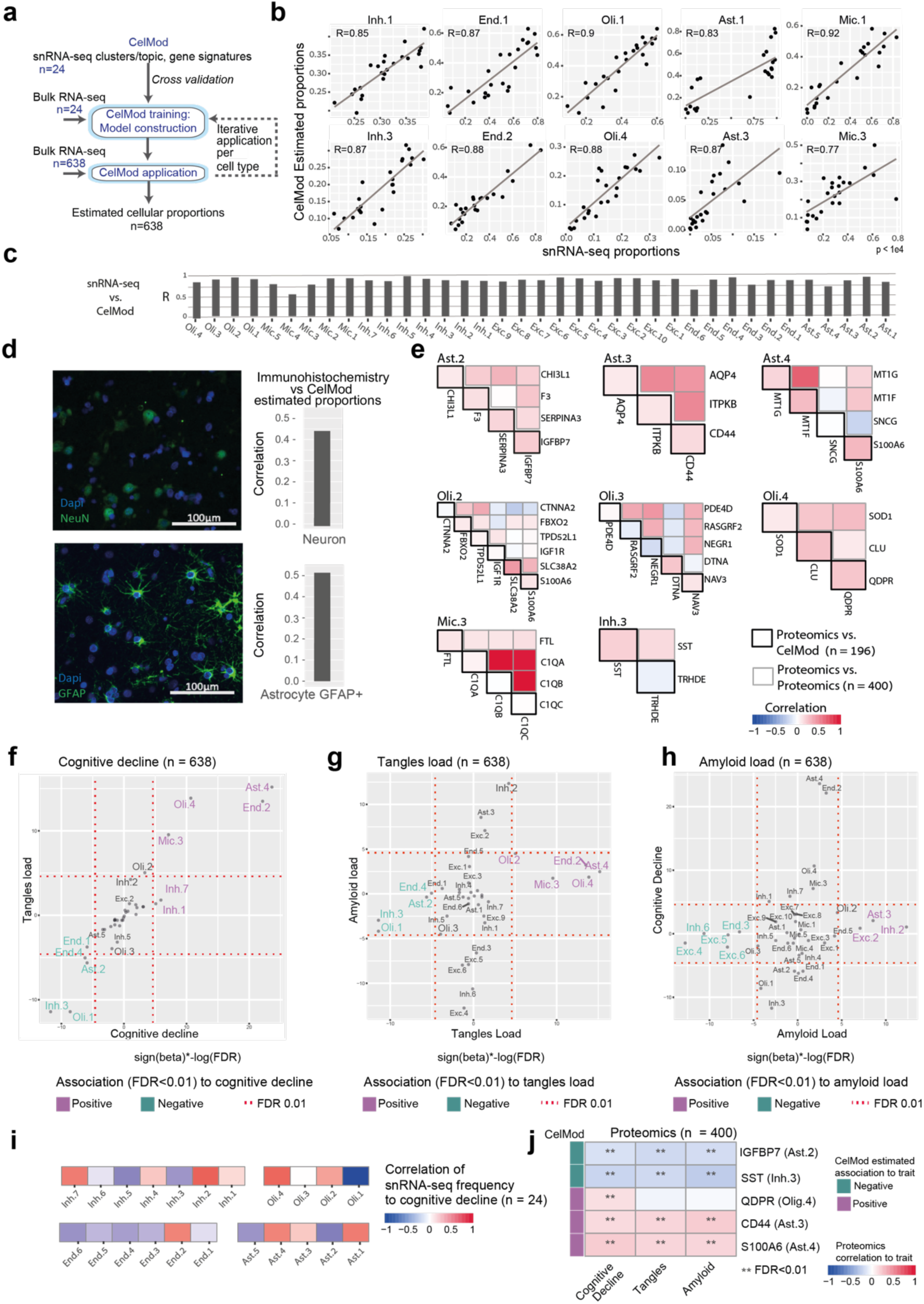
Proportions of cell subsets are associated with AD-traits in a cohort of 638 individuals. (**a**) Scheme of CelMod algorithm. Input: Cell types and cell subsets signatures and proportions across individuals derived from snRNA-seq samples (n=24). A two-step algorithm estimates cell subsets proportions in bulk RNA-seq samples (n=638), training on matching samples (n=24) using a four-fold cross-validation (Methods). **(b)** Estimated proportions of subsets by CelMod algorithm match snRNA-seq data. Scatter plots of the CelMod estimated proportions (Y-axis) compared to the snRNA-seq measured proportions (X-axis) across the 24 individuals for different subsets (cell type proportions and additional cell subsets are presented in **Extended Data Fig. 6a,b**). (**c**) Spearman correlation of the CelMod estimated proportions and the snRNA-seq measured proportions for each cell subset across the 24 individuals. (**d**) Cell type proportions measured by histology match CelMod estimates. Immunohistochemistry in DLPFC sections of 49 individuals (24 healthy, 24 declined), stained for marker for neurons (anti-NeuN, top) and reactive astrocytes (anti-GFAP, bottom). Left: Representative immunofluorescence images. DAPI, nuclei. Scale bar = 100 µm. Right: Pearson correlation coefficient of CelMod and immunofluorescence estimations of proportion (out of the total number of cells) for total neurons and for GFAP+ astrocyte populations (Astr.2, Astr.3). (**e**) Protein expression of signature genes of select cell subsets are correlated with CelMod estimates. Pairwise correlations of protein expression levels of subset signatures (blue-red color scale, gray squares, n=400 individuals), and correlations of protein expression to CelMod estimates (blue-red color scale, black squares, n=196 individuals, measured by both bulk RNA-seq and proteomics^58^). (**f-h**) Association of cell population to AD traits. Association scores for the CelMod estimated proportions across 638 individuals of all cell subsets (cell subtypes, states or topic models) to cognitive decline rate (f, X-axis), tangle burden (g, X-axis), β-amyloid burden (h, X-axis). Association score = −log(FDR)*sign(beta), from multivariable linear regression analysis corrected for age, sex and RIN values. Marking statistically significant subsets (FDR<0.01) with positive association (purple) or negative association (turquoise). (**i**) snRNA-seq empiric proportions of cellular subsets correlate with cognitive decline. Correlation (color scale) of proportions of cell subsets *within* each cell type to cognitive decline. Proportions are the fraction of nuclei within the subset out of the total number of nuclei within the cell type (for each of the 24 individuals with snRNa-seq profiles). Associations to additional AD traits in **Extended Data Fig. 6e**. (**j**) Protein levels of signature genes of selected subsets correlate to AD traits matching CelMod estimates. Correlation (color scale) of protein levels (rows) to cognitive decline rate, β-amyloid burden and tangles burden measured in n=400 individuals. ** = FDR < 0.01. Left bar: direction of association of the CelMod estimated proportion with the traits (purple: positive, turquoise: negative).

CelMod relies on a consensus of gene-wise regression models, with cross validation to estimate accuracy (**Methods, Fig. 4a**). Unlike most methods, CelMod is sensitive enough to detect the proportions of the different cell subsets (subtypes and states) *within* each broad cell class, and it can also deconvolve the relative contribution of expression programs (topics) in bulk profiles using program-specific weights (as opposed to proportions of discrete cell populations). We applied CelMod first at the broad cell class level and then separately within each class (**Fig. 4a**). CelMod identifies a large set of informative genes for each cell subset, ensuring that a small set of overlapping gene markers from different cell groups are not skewing the proportion estimates for broad cell classes as well as for subsets within each cell class. Ultimately, this resulted in inferred relative proportions of each of the 33 cell subsets and 4 oligodendrocyte topics in the bulk DLPFC RNA-seq data from each of the 638 ROSMAP participants (**Supplementary Table 4**).

CelMod accurately inferred proportions of snRNA-seq-derived cell types, cell subsets and programs (topics) as validated by comparing the inferred to the matched, empirically measured, proportions from snRNA-seq data using four-fold cross-validation: mean r=0.90, stdev=0.05 across cell types and mean r = 0.86, stdev = 0.09 across cell subsets *within* each cell type (**Fig. 4b-c, Extended Data Fig. 6a,b, Methods**). Of note, rare cell subsets have lower prediction accuracy, including the interferon associated Ast.5 and the redox associated Mic.3 (**Extended Data Fig. 6b**). CelMod showed higher prediction accuracy compared to previous deconvolution based methods^24–26^, which do not use cell proportion estimations explicitly in their training (**Methods, Extended Data Fig. 6c**). There was also agreement between the inferred proportions to matched proportions derived from immunofluorescence data on DLPFC tissue sections from the same brain region, yet in the contralateral fixed hemisphere^27^, in 49 ROSMAP participants with both bulk RNA-seq and histological data (**Methods**). Namely, we find good correlation for the proportion of neurons (NeuN+, r=0.40), and reactive astrocytes sub-population (reactive marker GFAP+, r=0.49) (**Fig. 4d**). We further validated the estimated CelMod proportions from proteomics measurements of key signature genes within key cell subsets across 196 individuals (with both bulk proteomics^28^ and bulk RNA-seq^1^ from the DLPFC), focusing on key cell subsets (Ast.2, Ast.3, Ast.4, Mic.3, Inh.3) and oligodendrocytes topics (Olig.2, Olig.3 and Olig.4) for which sufficient signature genes were measured in shotgun proteomic data (**Methods, Fig. 4e**). We first confirmed that the signature genes of each of the subsets/topics are co-expressed at the protein level, finding overall high pairwise correlations among protein levels (e.g. average R=0.22 and 0.51 for proteins within the Ast.2 and Ast.3 signatures, respectively) (**Fig. 4e**). Next, we validated that the expression levels of the signature proteins across individuals are correlated with the CelMod estimated proportions of the relevant cell subsets (**Fig. 4e**). Despite the reported differences between brain RNA and protein levels^29^, we found correlations between protein levels and CelMod proportions for most cell subsets (e.g. R = 0.23,0.17,0.3,0.24,0.2, for IGFBP7 and Ast.2, CD44 and Ast3, S100A6 and Ast.4, QDPR with Oli.4, SST with Inh.3 frequency, respectively) and lower correlations for a few others (e.g. R=0.06 for C1QA with Mic.3 frequency, and R=0.01 for SNCG with Ast.4 signature). Notably, bulk protein and bulk RNA levels of C1QA are lowly correlated (R=0.14), and they are anti-correlated for SNCG (R=−0.4) (**Fig. 4e**). Overall, some cell subsets appear to be inferred more accurately than others; nonetheless, we assess all inferred subsets in the next steps of the analysis, as poor inference will tend to bias analyses of affected cell subtypes towards the null and would be less likely to create false positive associations.

### Cell population frequencies are associated with AD-pathologies and cognitive decline

With this validated set of inferred cell subset proportions in 638 individuals, we next turned to exploring the association of cell subset proportions with AD-related traits available from the deep ante-mortem and post-mortem characterization of ROSMAP participants^7–9^. We prioritized three outcomes that capture distinct critical aspects of AD: quantitative measures of (1) β-amyloid and (2) tau burden, which are the two defining pathologic characteristics of AD, as well as (3) the slope of aging-related cognitive decline over up to 20 years before death. This last measure captures the progressive cognitive impairment that leads to dementia. β-amyloid generally accumulates earlier than tau pathology, but, while both pathologic features are found in the DLPFC of most ROSMAP participants, tau burden is more closely associated with cognitive impairment and dementia^30,31^ (**Extended Data Fig. 6d**). We thus tested for association of these three AD-related traits with the inferred cellular proportions (*within* each broad cell class) using linear regression (with correction for age, RIN score and sex, **Methods, Supplementary Table 5, Fig. 4f-h**).

We revealed significant associations (FDR<0.01) between the proportions of neuronal and glial cell subsets (as measured *within* each cell class, **Methods**) and the pathologic and/or cognitive traits (**Fig. 4f-h**). Most cell subsets and topics associated with tau pathology also showed significant associations to cognitive decline (FDR<0.01, **Fig. 4h**); these include a positive association with cognitive decline and tau pathology for Oli.4, Ast.4, Mic.3, End.2, and as well as negative associations for Oli.1, Inh.3, End.4 and Ast.2. Few subsets were only associated with cognitive decline: End.1 negatively and Inh.1/7 positively (FDR <0.01), while Oli.2 positively associated with tau pathology (FDR<0.01) and only marginally associated with cognitive decline (FDR < 0.036). Conversely, although variation in proportions of several cell subsets was significantly associated with β-amyloid pathology (FDR<0.01), none of these were significantly associated with cognitive decline. These included negative associations with subtypes of glutamatergic neurons Exc.4/5/6 (layer 4-5 pyramidal neurons), and inhibitory neuron subtype Inh.6 (expressing PTPRK, inferred to be found predominantly in layer 4) and to an endothelial subset (End.3), and positive associations with Ast.3 (GFAP+CD44+), Exc.2, and Inh.2 (**Fig. 4h**). This overall finding is consistent with the stronger association between tau (vs. β-amyloid) pathology and cognitive decline^30,31^. β-amyloid pathology, while correlated with Tau pathology (r=0.48, **Extended Data Fig. 6d**), each have a range of strong cellular associations (FDR<0.01), only some of which are also marginally correlated with the other pathology (FDR<0.05, e.g. Oli.1, End.2, Ast.4 and Inh.3). Thus, the two pathologies have a largely distinct set of associations with cell subtypes/states, consistent with our earlier report of distinct microglial transcriptional programs being associated with amyloid and tau pathology^27^.

The inferred associations with cognitive decline and tau-pathology showed that oligodendrocyte topic Oli.1 (SVEP1+) appears to be more prominent in non-impaired individuals while topic Oli.4 (QDPR+) is enriched in individuals with cognitive decline. In parallel, the relative proportion of inhibitory neuronal subtype Inh.3 (SST+) is higher in healthy individuals, in contrast to inhibitory subtype Inh.1 (PV+), which is relatively higher in individuals with cognitive decline (**Fig. 4f-h**). This suggests a potential high vulnerability of SST+ GABAergic neurons in AD. Additional subsets positively associated with cognitive decline include End.2 and Ast.4 which are enriched for genes related to stress response pathways such as oxidative stress (**Fig. 3f,i**).

Although our sample size of single nucleus data has limited statistical power compared to the larger set of inferred cell type proportions, we still found a clear trend of positive or negative correlations between the proportion of certain cell subsets in the snRNA-seq data (n=24, *within* each broad cell class) and cognitive decline, β-amyloid and tau burden (**Fig. 4i, Extended Data Fig. 6e**), which matches our findings from the CelMod estimations in 638 samples. For example, the relative proportions of the Olig.1 topic, of Ast.2 and End.4 subsets, and of the Inh.3 subtype are both negatively correlated with cognitive decline (*r*=−0.50, −0.318, −0.24, −0.08 and −0.26 respectively), while Olig.4, Ast.4, End.2 and Inh.1/7 proportions are positively correlated (r=0.38, 0.31, 0.31, 0.09 and 0.33 respectively) (**Fig. 4i**).

We further validated the association between cell subtypes/subsets and the AD traits by additional bulk RNA-seq profiles and proteomic measurements. Appling the inferred CelMod model to an independent set of 106 bulk RNA-seq profiles from a different cohort (from the Mount Sinai Brain Bank (MSBB^32^), **Methods**), yielded matching cellular association to both cognitive decline as well as to Tau burden for most subsets (except for Ast.2, End.4, and Mic.5 subsets, **Extended Data Fig. 6f**). In parallel, we used proteomic measurements for selected proteins as markers for key prioritized subsets in 400 ROSMAP individuals^28^. Consistent with our RNA based analysis, the SST protein (Inh.3 marker) and IGFBP7 (Ast.2 marker) are anti-correlated with cognitive decline and tau burden (p<0.01, **Fig. 4j**). In addition, we validated the positive association between Oli.4 and cognitive decline using the marker QDPR protein levels as well as the association of Ast.4 with cognitive decline, tau and amyloid using the marker S100A6 protein levels (p<0.01, **Fig. 4j**). Finally, with the CD44 protein levels, we validated the positive association of amyloid burden with Ast.3, and we found that, at the protein level, CD44 also positively correlated with cognitive decline and tangles (p<0.01, **Fig. 4j**).

### Co-variation structure of cell subsets proportions across 638 individuals uncovered multicellular communities in the aging neocortex

Assessing the extent of inter-individual variation in cellular composition across the 24 participants with snRNAseq data revealed differences *within* each of the cell classes separately, uncovering variability in the frequencies of cell subsets across individuals, which was beyond the association of specific cell subsets to the archetype of individuals (**Fig. 5a**). To assess the robustness of these differences in frequency of cell subsets across individuals, we used the CelMod-inferred proportions of each of the 33 cell subsets and 4 oligodendrocyte topics from the bulk DLPFC RNA-seq across 638 participants (**Supplementary Table 4**). We found co-variation in cell subset frequency (*within* each cell type) across these 638 participants, similar to what we initially observed for the 24 participants (**Fig. 5a-b**).

**Figure 5.**
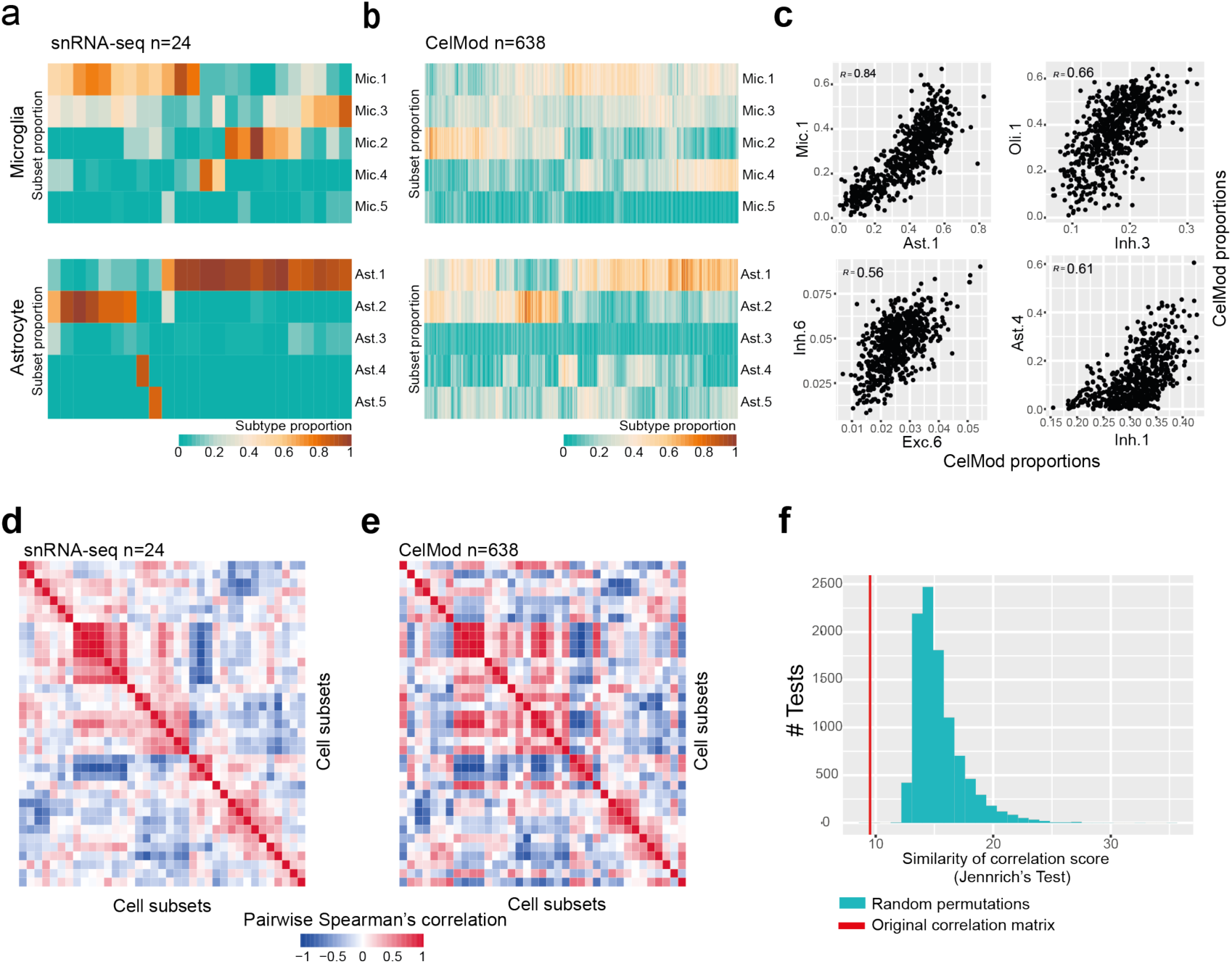
Co-variation structure of cell subsets proportions across 638 individuals. (**a-b**) Proportions of cell subsets across individuals in astrocytes and microglia. The frequency (out of total cells in the class, color scale) of each cell subset (columns) in each individual (rows): from snRNA-seq (n=24, in a) or CelMod estimations (n=638, in b). The cell subsets in (b) are ordered as in (a). (**c**) Scatter plots of selected pairs of cell subsets from different cell classes, showing high correlations between proportions of subsets (n = 638). (**d-e**) Coordinated changes in proportions of cell states and subtypes across individuals. A heatmap of the pairwise Spearman’s correlation coefficient of the proportions of all cell states and subtypes across 24 individuals (snRNA-seq measurements, d) and coordinately in 638 individuals (CelMod estimated proportions, e). Exposing a structure of mixed correlated and anti-correlated cellular subsets of mixed cell types. (**f**) CelMod estimated pairwise correlations of cellular populations (n=638 individuals) matches the snRNA-seq measurements (n=24 individuals). Similarity between the two correlation matrices (in d and e) is statistically significant (p-value<0.001, by permutation test, **Methods**). Histogram of the distribution of similarity scores (Jennrich’s test ^60^) of correlation matrices in 10,000 random permutations of the cellular frequencies independently per cell type. Red, similarity score of the true matrices in d and e.

We next asked how the variation across all the individuals in the proportion of each cell subsets was correlated with variation in other cell subsets. To this end, we calculated the Spearman correlation coefficient for each pairwise combination of cell subset or topic proportions (where proportions are defined *within* a cell class, **Fig. 5c**), uncovering a co-variation structure of cell subsets from multiple cell types captured by hierarchical clustering of these pairwise correlations (**Fig. 5d-e, Methods**). The cell subsets exhibited significantly similar co-variation structures across both the 24 participants with empirically determined proportions (**Fig. 5d**) and across the 638 participants with inferred proportions (**Fig. 5e**), as assessed by a permutation test (**Fig. 5f**, p-value<0.001, **Methods**). We confirmed the cellular architecture of the neocortex in bulk RNA-seq profiles of 106 individuals from the independent MSBB cohort for which we inferred cell subset proportions using CelMod (**Methods, Extended Data Fig. 7a**).

Correlated proportions of cell subsets across individuals suggest the existence of distinct multi-cellular communities in the aging human brain. To define those, we built a network of cell subsets, where we connected each pair of subsets (nodes) that are significantly correlated or anti-correlated (signed edges) (R>0.4 or R<-0.4, with p<0.05, **Methods, Fig. 6a,b**). These networks identified an underlying structure of cellular communities (connected components of positively correlated cell subsets) composed of cell subsets across multiple neuronal, glial and endothelial cell types, along with strongly opposing communities (negatively correlated cell subsets between communities) (**Fig. 6b**). A similar architecture was detected in the 24 snRNA-seq derived network (**Extended Data Fig. 7b**).

**Figure 6.**
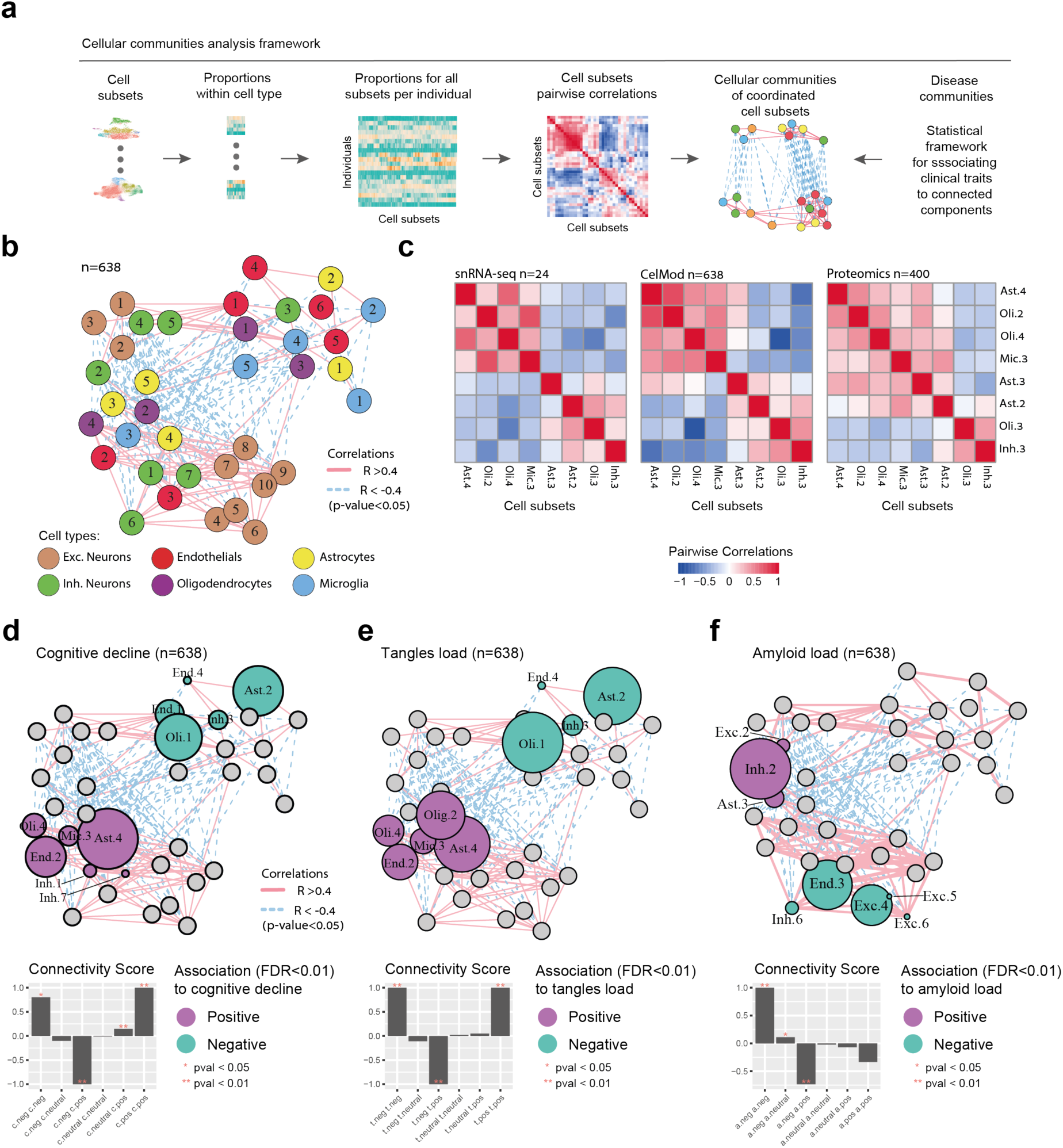
Multi-cellular communities exist in the aging DLPFC brain region. (**a**) Scheme of computational framework for estimating multi-cellular communities: Proportions of cell subsets across individuals within each cell type are calculated, combined, and pairwise correlations between all cellular subsets are computed. A multi-cellular network is derived from the pairwise correlations, associated with AD-traits by statistical analysis, and connected components are annotated as cellular communities (**Methods**). (**b**) A network of cellular subsets reveals coordinated variation across individuals in multiple cell types. Network of coordinated and anti-coordinated cell subsets (nodes). Edges between pairs of subsets with significantly correlated proportions across individuals (r>0.4, p-value < 0.05, solid red line) or anti-correlated (r<-0.4, dashed blue line) based on CelMod proportions (n=638). snRNAseq based network in **Extended Data Fig. 7b**). Nodes are colored by the cell type and numbered by the subset as in Fig. 2a **and Fig. 3a,d,h**). (**c**) Correlation pattern of proteomics^28^ expression of signature genes across cell subsets match CelMod estimates. For selected cell subsets, pairwise correlations of: snRNA-seq proportions (n=24, left), CelMod proportions (n=638, middle), and average protein expression level of signature genes (n=400, right). (**d-f)**. Cellular communities are linked to AD associated traits. Cellular network (as in **b**), of coordinated and anti-coordinated cell subsets (nodes), colored by the associations (multivariable linear regression, FDR<0.01) with AD traits (purple: positive, green: negative association, grey: non-significant) for: cognitive decline (**d**), tangles burden (**e**), and β-amyloid burden (**f**). Bottom: Each bar represents the connectivity score ((#positive edges - #negative edges)/potential edges) between groups of cells according to their association to each trait, showing that cell subsets associated with AD-traits are highly connected in the network. Statistical significance for the connectivity score calculated based on random permutations (Mathods): *<0.05, **<0.01.

We validated, at the protein level, the coordinated levels of cell subsets across individuals for subsets of different cell types using bulk proteomic data. Consistent with the two transcriptomic networks (CelMod estimated n=638, and snRNA-seq measured n=24), there were significant correlations between the protein level of signature genes of various cell subsets: a positive correlation among signatures of Ast.4, Oli.2, Oli.4 and Mic.3 (average R=0.31), and, separately, between signatures of Oli.3 and Inh.3 (R=0.41), along with an anti-correlation between the two sets (average R=−0.2). Notably, the are some differences between the RNA and protein-derived results: in correspondence with the RNA based analysis, the Ast.2 signature correlated with Inh.3 signature and Ast.3 signature correlated with Oli.1 but an additional correlation of both Ast.2 and Ast.3 signatures with Mic.3 emerged only in the proteomic data (**Fig. 6c**). This highlights the need for multi-omic data integration to better capture inter-cellular relationships.

### A neuro-glial-endothelial multicellular community is associated with cognitive decline

We next asked whether the changes in the composition of cell subsets associated with AD (**Fig. 4g-h**) reflect multiple independent effects that contribute to cognitive decline, or a coordinated set of changes that is captured by our cellular networks, which is more likely given the dependency structure we uncovered (**Fig. 6b**).

Cell subsets associated with AD related traits formed connected communities in our networks (**Methods, Fig. 6d-f**). In particular, the cognitive decline-associated cell subsets segregated into two anti-correlated communities, each composed of different highly interconnected neuro-glial-endothelial cell subsets: (1) the *cognitive decline community* consisting of cell subsets whose proportions were associated with more rapid cognitive decline (*p-value* < 0.01 compared to random, Methods) as well as greater tau pathology burden (*p-value* < 0.01). This community includes the Olig.4 signature, Astr.4, Micr.3, End.2, and GABAergic PV+ neuronal subtype Inh.1 and Inh.7 subtypes (**Fig. 4f,g, Fig. 6c,d**). On the other hand, (2) the *cognition non-impaired community* consisted of cell subsets whose proportions were significantly higher in individuals with little or no cognitive decline and low Tau pathology (*p-value* < 0.01), including the Olig.1 signature, the GABAergic SST+ neuron, Inh.3, Astr.2 and End.1 subsets (**Fig. 4f,g, Fig. 6d,e**). For β-amyloid burden, the *Amyloid low community,* included neuronal cell subsets that are negatively associated with β-amyloid burden, specifically, glutamatergic neuronal subtypes Exc.3-6 and Inh.6 (p<0.001, **Fig. 4h, Fig. 6f**). To statistically test the connected structure of multi-cellular communities, we used a random permutation test and applied a connectivity score to each community, showing that cell subsets associated with cognitive decline, tau burden, and low amyloid burden, captured significantly (p<0.01) connected components in the network. The connections between reciprocal communities, such as the cognitive decline vs. cognition non-impaired were found to be significantly connected by negative edges (p<0.01). We therefore identified several sets of coordinated cellular responses that are associated with distinct aspects of AD, with a tight neuro-glial community associated with accumulation of both proteinopathies as well as cognitive decline.

Notably, excitatory neuronal subsets largely formed two opposing, independent connected components in the graph: one consisting of upper cortical layer neuronal subsets (layers 2-3) and the other of lower layer neuronal subsets (cortical layers 4-6, **Fig. 6b**). Given the intrinsic association between pyramidal neuron subsets and cortical layers (**Fig. 2b**), we cannot exclude the possibility that this partition is driven by a dissection bias of the cortical samples, yet as this dependency structure was captured in both the single and bulk RNA-sequencing profiles (**Fig. 5d,e** and **Extended Data Fig. 7a**), this explanation is less likely. Notably, other cell subsets (inhibitory, endothelial and glial subsets) were found to be independent of the cortical layer proportions and thus we can conclude their proportions are not affected by a dissection-related bias (**Methods, Extended Data Fig. 7c**). Of note, previous work found increased neuronal vulnerability associated with AD pathology in RORB+ pyramidal neurons positioned at middle-deep cortical layers^10^, consistent with the negative association we observed between amyloid pathology and our excitatory pyramidal neuron subsets Ex.4, Ex.5 and Ex.6 (**Fig. 4h**) positioned at layers 4-5 of the neocortex (**Fig. 2**) and the Ex.4 and Ex.5 subtypes both express RORB (**Extended Data Fig. 2a**).

### Distinct pathways and ligand-receptor pairs identified in the cognitive decline and tau pathology associated cellular communities

The existence of co-occurring cell subsets we termed *cellular communities* (**Fig. 6**) suggests that certain cellular populations, may reflect shared functionalities as well as distinct signaling patterns between cell subsets *within* each community^33,34^, and may be engaged in a common disease-related process. We thus searched for shared enriched pathways as well as for ligand-receptor interactions connecting cell subsets within and between communities, focusing on the two opposing communities associated with cognitive trajectory, the *cognition non-impaired* community (Inh.3, Oli.1, Ast.2, End.1, End.4) and the *cognitive decline* community (Inh.1, Inh.7, Oli.4, Ast.3, Mic.3, End.2)

Examining the differentially expressed pathways across each cell subsets (compared to all other subsets of the same cell type) within the *cognition non-impaired* community and within the *cognitive decline* community, we found 14 and 275 overlapping pathways for at least 3 of the cell subsets within each community, respectively (hypergeometric test, qvalue<0.05). Interestingly, the *cognitive decline community* cell subsets shared pathways related to known risk factors associated with AD, including response to oxidative stress, hypoxic stress, and DNA damage. They also shared an up-regulation of oxidative phosphorylation and neurodegenerative diseases associated genes (including AD) as well as translation regulation (qvalue<0.05, **Fig. 7a**). On the other hand, the cell subsets within the *cognition non-impaired community* shared pathways related to axon development, cell-cell junction and synapses related pathways (qvalue<0.05, **Extended Data Fig. 7d**).

**Figure 7.**
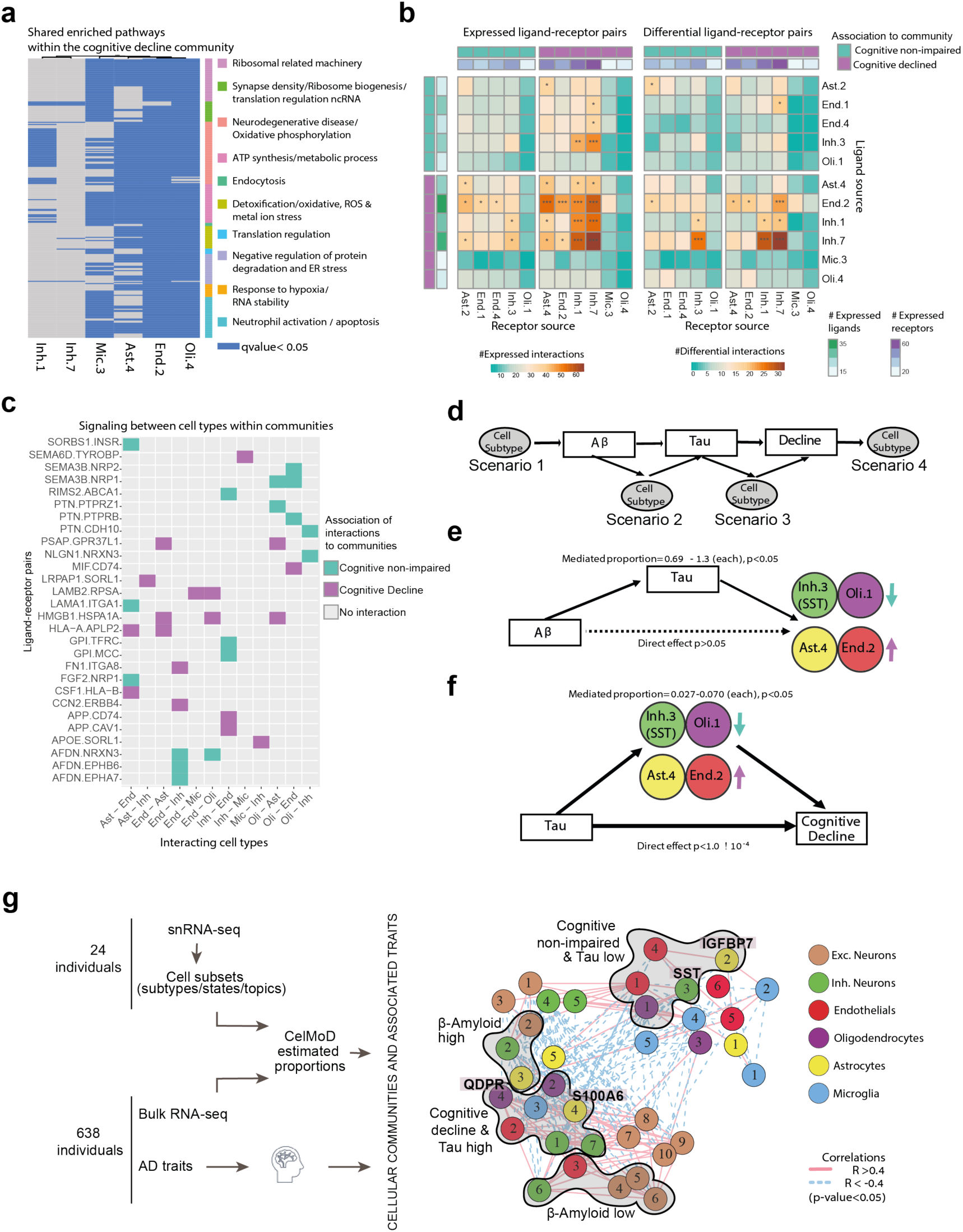
The cellular environment of the AD and the cognitively non-impaired brains. (**a)** Shared pathways within the cognitive decline community. Enriched pathways (hypergeometric test, qvalue<0.05) in up-regulated genes within each cell subset, shared between at least three subsets within the cognitive decline community. Pathways are clustered by shared genes (color bar, Methods). Shared pathways for the non-impaired community in **Extended Data Fig. 7d**. (**b**) Cell subsets positively associated with cognitive decline have an increased expression of ligand-receptor pairs compared to the negatively associated subsets. For each pair of cell subsets showing the numbers (color scale) of ligand-receptor pairs (LRP, row: ligand, column: receptor) that are expressed (**left**) or differentially expressed in at least one of the subsets (**right**). Top and side bar marking: subsets positively associated with cognitive decline (purple) or negatively (turquoise), and the general number of expressed LRPs (color scale). (**c**) Examples of community-specific LRPs. For each LRP, marking the association to each pair of cell subsets with the *cognitive declined* community (purple) or the *cognitive non-impaired* community (turquise) or no-association (grey). Associating only LRPs in which both the ligand and the receptor are positively differentially expressed in the relevant pair of cell subsets. (**d**) A scheme of our underlying assumptions and the four possible scenarios of the cell subset causal relationship with AD pathology and cognitive decline assessed by mediation analysis. (**e**) Mediation analysis results showing tau pathology burden is predicted to be upstream of changes in proportion of Inh.3, Olig.1, Ast.4, and End.2. Colored by cell type, the arrow indicates the direction of change in proportion in association with cognitive decline and tangles burden. Full results in **Extended Data Fig. 8c**. (**f**) Mediation analysis results showing partial effect of changes in proportion of Inh.3, Olig.1, Ast.4, and End.2 on cognitive decline independent of tau pathology burden. Colors and arrows as in e). Full results in **Extended Data Fig. 8d**. (**g**) A scheme of our proposed model of multi-cellular communities of the aging DLPFC brain region and their associations with AD traits. Cellular networks (as in **a**), nodes colored by the community assignments. The significantly enriched associations to AD-traits (hypergeometric p-value) are marked next to the graph.

We searched for ligand-receptor pairs (LRPs)^35,36^ that putatively connect different cell subtypes within and between the *cognitive decline community* and the *cognition non-impaired community* (**Methods**). We identified a total of 298 unique *expressed* LRPs (13.8% of all possible LRPs, p<0.05, inferred by a permutation test by CellPhoneDB^37^, **Methods**), connecting one cell subset expressing the ligand with another expressing the receptor (**Methods**) (**Fig. 7b**). The number of expressed LRPs was higher between cell subsets within the cognitive decline community (1,005 LRPs) as opposed to the cognition non-impaired community (458 LRPs) or between these two communities (681 and 684) (**Fig. 7b**).

Next, we searched for *community-specific* LRPs, defined as expressed LRPs in which at least the ligand or the receptor were differentially expressed within a cell type between the two communities (FDR<0.01, **Fig. 7b-c, Extended Data Fig. 8a,b, Methods**). We identified 383 *cognitive decline* community-specific LRPs, such as the Laminin Subunit Beta 2 (LAMB2) expressed by End.2 and Laminin Receptor 1 (RPSA) expressed by Mic.3 and Oli.4 respectively (FDR<0.01, **Fig. 7c**). Within the *cognition non-impaired* community we found 181 community-specific LRPs, such as the Fibroblast Growth Factor 2 (FGF2) expressed by Ast.2 and the receptor Neuropilin 1 (NRP1)^38^ found in End.1, FDR<0.01, **Fig. 7c**). We further identified pairs of cell subsets statistically significantly enriched in community-specific LRPs (permutation test, p<0.01, **Methods, Fig. 7b**), and found that most such pairs of cell subsets were within the *cognitive decline community* (**Fig. 7a,b**).

### The proportion of certain DLPFC cell subsets partially mediate the association between tau pathology and cognitive decline

While our autopsy-based cross-sectional data cannot formally determine the causal chain of events leading to AD, modeling can propose a most likely scenario. To test whether cell proportion changes may mediate the association between AD pathology and cognitive decline, we analyzed the cell subsets associated with our quantitative AD-related traits in a causal modeling framework. We built our models based on the widely assumed sequence of AD progression, that starts with Aβ accumulation, followed by tau accumulation with subsequent neurodegeneration and cognitive decline^39,40^. This is also consistent with our data, in which no cell subset was directly associated with both amyloid load and cognitive decline without being associated with tau^41,42^ (**Fig. 4f,g**). Our mediation modeling thus aimed to align cell subtype proportions along this sequence of AD pathophysiology (**Fig 7d**).

First, we considered the relation between Aβ, Tau, and cell subtype proportions. We focused on the 6 tau-associated cell subsets that were also associated with Aβ at a nominal level (p<0.05, but FDR>0.017, Inh.2, Oli.1, Oli.2, Ast.4, End.2, Inh.3(SST)). We tested three possible scenarios based on the known direction of effect from Aβ to tau (**Fig 7d,e**), where cell subset proportion changes are either (**1**) upstream drivers of Aβ levels; (**2**) mediators, downstream of Aβ but upstream of tau; (**3**) downstream of tau. In each scenario, we tested the change in association between two of the variables once the third is added as a mediator. Four out of six cell subsets tested (Ast.4, End.2, Inh.3 and Oli.) were precited to be downstream of Tau, as tau burden mediated most of the amyloid– cell subtype associations (proportion mediated > 69%), leaving no significant direct amyloid effect on cell subset proportions (p>0.05) (**Fig. 7e, Extended Data Fig. 8c**). While for Oli.2, a marginal direct amyloid – cell subset proportion effect was also observed (p=0.05), suggesting that Oli.2 proportion might be affected by both amyloid and tau (**Extended Data Fig. 8c**). For Inh.2, we predict that it might be upstream of tau since Inh.2-tau association almost completely attenuated after adjusting for Aβ (**Extended Data Fig 8c**), suggesting that changes in the frequency of these inhibitory neurons may be occurring at an earlier stage, amyloid-centric stage of AD.

Next, having positioned cell proportion changes for Ast.4, End.2, Inh.3 (SST+) and Oli.1 as being likely downstream of tau accumulation, we performed a second mediation analysis to align these cell subsets along the sequence of events leading to cognitive decline (**Fig. 7d,f**). Our analysis positioned all four cell subsets upstream of cognitive decline, as each was found to partially mediated the tau-cognitive decline relationship (2.7 – 7.0 % of total effect), while tau retained a strong direct effect on cognitive decline (>90% of total effect) (**Fig. 7f, Extended Data Fig. 8d**). When the proportions of the four cell subsets were considered simultaneously, they explained 7.4% of the tau – cognition association. Thus, the changes in the frequency of these four cell subtypes explains a small but meaningful fraction of the effect of tau proteinopathy on cognitive decline, laying the foundation for the development of new hypotheses to preserve cognition.

## DISCUSSION

In this study, we constructed a high-resolution cellular map of the DLPFC, which enabled the characterization of the diversity of neuronal, glial and endothelial cell subsets and expression programs at high resolution in order to extract new insights about intra- and inter-individual diversity in the aging brain and about coordinated multi-cellular *communities* associated with cognitive impairment and hallmark AD pathologies (**Fig. 1**). Our optimized snRNA-seq protocol and analytic pipelines uncovered a large diversity of cellular populations in the aging neocortex in 24 individuals (**Fig. 2 and 3**, compared to previous reports^2,5^, **Extended Data Fig. 4**). Further, using a new method (CelMod), we integrated this detailed cellular map with bulk RNA-seq data from the same brain region, to estimate the proportions of the major cell classes and cell subsets in 638 participants in the same aging cohorts (ROSMAP) and validated in 106 participants from an independent cohort (MSBB) (**Fig. 4, Extended Data Fig. 6**). The general approach can be readily applied to other matched tissue-level datasets. The inferred cellular composition data was statistically powered for disease association analyses, which uncovered new observations such as a striking decrease in SST+ (Inh.3) GABAergic neuron proportions, suggesting that SST neurons may be more vulnerable to tau pathology, validated by proteomics of the same brain region in 400 individuals (**Fig. 4 and 6**). Such changes were not reported in previous studies^2,5^, perhaps because they had profiled a lower total number of neurons. Next, we applied a new computational framework to expand the analysis from a cell-type centric view to a multi-cellular view of cellular environments and organization, which led to the discovery of distinct *cellular communities* that we linked to AD associated traits (**Fig. 6**).

In microglia, we mapped our snRNA-seq-based clusters to those found in single-cell analysis of live human microglia purified from fresh autopsy and surgically resected tissue ^6^, finding a good match with only a few differences in subtype composition between the two experimental approaches, addressing an important concern raised in an earlier study^12^. As a result, although there are important differences relating to the quantity, quality and nature of single nucleus- and cell-derived RNA-seq data of microglia, no microglial subtype seems to be missing in either dataset. This suggests that these data could be integrated in future studies. Notably, while we cannot completely exclude technical artifacts from *ex vivo* manipulations of cells and nuclei, the similarity in microglia diversity captured by both live cells and frozen nuclei substantially reduces the likelihood that such artifacts have a strong effect on results.

Oligodendrocytes emerge as an interesting cell type for further evaluation. Oligodendrocytes were notable both for their association to AD and because they highlight the need to accommodate the unique characteristics of each cell type as analytic methods are selected (**Fig. 3j, 4f-g**): one method may not be optimal for all cell subtypes in a tissue specimen. We represented the heterogeneity within these cells utilizing a continuous approach (topic models ^18–21^) to capture expression programs instead of discrete cell clusters. One such topic (Olig.4) is very strongly associated with tau pathology and cognitive decline, while another (Olig.1) is prevalent in cognitively non-impaired brains, suggesting a clear oligodendrocytic signature associated with AD (shown by RNA signatures and protein markers expression, **Fig. 4, 6c**).

A major innovation of our study is the definition of cellular *communities* defined by correlated changes in the frequency of different cell subsets across individuals (defined within each cell class, **Fig. 6 and 7g**). The correlation structure initially observed in our snRNA-seq data from 24 individuals (**Fig. 5a,e**) was reproduced in the 638 ROSMAP participants who have inferred cell type proportions (**Fig. 5b,e**). Within the cognition non-impaired- and cognitive declined-communities we found shared pathways across the different cell subsets separately for each community, highlighting pathways of known risk factors for AD that were found to be shared within the declined community (**Fig. 7a, Extended Data Fig. 7d**). We also found expression of known ligand-receptor pairs enriched within subsets of the declined community (**Fig. 7b-c**). These results fit within our conceptual understanding that AD is a distributed pathophysiologic process involving multiple interacting cell types. Spatial transcriptomics methods will help to resolve whether these communities represent groups of co-located cells or distributed communities responding to an underlying, shared signal.

One strength of our analyses is their foundation of ROSMAP participants that have detailed, quantitative clinicopathologic measures that enable us to resolve “AD-related changes” related to β-amyloid or tau pathology from those involved in cognitive decline, the ultimate therapeutic target. This is nicely illustrated by a broad set of cell subset changes with β-amyloid that are not associated with cognitive decline and can therefore be deprioritized. These analyses also show, consistent with many earlier analyses, that the burden of tau pathology is more closely related to cognitive decline than the burden of β-amyloid. This leads us to prioritize those cell populations that, in our mediation modeling, appear to contribute to the consequences of tau proteinopathy: we implicate a clear set of 4 different cell subtypes whose changes in frequencies lead to cognitive decline as a result of tau proteinopathy. Thus, these cell subtypes become key targets for therapeutic development efforts.

Our study does have limitations: while we have statistically robust results thanks to CelMod estimates in bulk RNA-seq from 638 individuals, our reference snRNAseq data is from 24 participants and likely did not capture the full diversity of cell population in aging brains, especially in more challenging, less frequent cell populations such as microglia, OPCs and pericytes; validation of our results in large-scale snRNAseq studies is necessary to ensure that they are robust. Specifically, we have a limited ability to reliably estimate the abundance of rare cell subsets such as Mic.3, End.4 and Ast.5. Further, we have validated the observed cellular architecture and association to traits using histology, RNA and proteomics data; however, the associations are limited by the detection limits of each method and by differences between RNA and protein expression levels. It is likely that larger-scale snRNA-seq studies will enable us to better resolve additional cell subtypes and transcriptional programs and to uncover additional, lower frequency cell subtypes that may be important in AD. Higher resolution spatial maps will help to resolve whether these communities represent groups of co-located cells or distributed communities responding to an underlying, shared signal. Finally, because profiling post-mortem brain tissue is by definition a cross-sectional study, we have to infer the temporal links between cellular communities, appearance of pathology, and cognitive symptoms. Our mediation analysis suggested that the strong known link between tau pathology and cognitive decline is mediated in part through the cell type proportion changes that we observed, specifically through an oligodendrocyte state changes and differences in SST+ interneuron proportions (**Fig. 7e,f**). Thus, we propose a set of hypotheses that can now be tested in longitudinal samples or model systems.

Overall, our work highlights the importance of a unified, cross-cell type view of the cellular ecosystems of the brain, beyond a cell-type-specific focus, in the study of AD and other complex neurodegenerative disorders. Embracing the complexity of this heterogeneous parenchymal tissue, network approaches can uncover new insights, such as key members of each cellular community that are involved in different aspects of the aging brain.

## Methods

### Experimental design

Data were derived from subjects enrolled in two clinical-pathologic cohort studies of aging and dementia, the Religious Orders Study (ROS) or the Rush Memory and Aging Project (MAP), collectively referred to as ROSMAP. All participants are not diagnosed with dementia at enrolment, have annual clinical evaluations and agree in advance for brain donation at death. At death, the brains undergo a quantitative neuropathologic assessment, and the rate of cognitive decline is calculated from longitudinal cognitive measures that include up to 20 yearly evaluations^7–9^. For this study, we used data from 24 individuals (12 males and 12 females) chosen to represent the range of pathologic and clinical diagnoses of AD dementia (at the time of death), divided to four groups (**Supplementary Table 1**): (1) a <u>reference group</u> of cognitively non-impaired individuals (cAD =1 in ROSMAP) with minimal AD pathology (pathoAD =0 in ROSMAP), (2) a <u>resilient group</u> of cognitively non-impaired individuals (cAD=1) with a pathologic diagnosis of AD (pathoAD=1), (3) an <u>AD group</u> who fulfill diagnoses for both clinical AD dementia and pathologic AD (cAD=4 and pathoAD=1), and (4) a <u>clinical-AD group</u> of individuals diagnosed with clinical AD dementia but showed only minimal AD pathology upon post-mortem characterization (cAD=4 and pathoAD=0). We included only samples that had RIN>5 and with post-mortem interval (PMI) < 24, and samples that had bulk RNA-sequencing in a previous study ^1^ as well as whole genome sequencing data^39–42^.

### Experimental design: AD traits in the ROS/MAP cohorts

The pathologies were collected as part of the ROS/MAP cohorts (previously described in details^43–46^). We used the following traits in the poise of samples and in the association analysis: ***Cognitive Decline.*** Uniform structured clinical evaluations, including a comprehensive cognitive assessment, are administered annually to the ROS and MAP participants. The ROS and MAP methods of assessing cognition have been extensively summarized in previous publications^9,44,47–49^. Scores from 17 cognitive performance tests common in both studies were used to obtain a summary measure for global cognition as well as measures for five cognitive domains of episodic memory, visuospatial ability, perceptual speed, semantic memory, and working memory. The **summary measure for global cognition** is calculated by averaging the standardized scores of the 17 tests, and the summary measure for each domain is calculated similarly by averaging the standardized scores of the tests specific to that domain. To obtain a measurement of cognitive decline, the annual global cognitive scores are modeled longitudinally with a mixed effects model, adjusting for age, sex and education, providing person specific random slopes of decline (which we refer to as ***cognitive decline***). The random slope of each subject captures the individual rate of cognitive decline after adjusting for age, sex, and education. Further details of the statistical methodology have been previously described^50^. **Clinical diagnosis of AD at the time of death.** Annual clinical diagnosis of AD dementia follows the recommendation of the joint working group of the National Institute of Neurological and Communicative Disorders and Stroke and the AD and Related Disorders Association ^51^. The diagnosis requires a history of cognitive decline and evidence of impairment in memory and at least one other cognitive domain. After a participant had died, a neurologist specializing in dementia reviews all available clinical information and provides a summary opinion with regards to the most likely clinical diagnosis at the time of death. The summary diagnosis was blinded to all neuropathologic data, and case conference are held for consensus as necessary ^52^. AD dementia includes persons with probable or possible AD dementia, i.e. AD dementia with or without comorbid conditions that may be affecting cognition. **A pathologic diagnosis of AD** was determined by a board-certified neuropathologist blinded to age and all clinical data and using modified Bielschowsky silver stained 6 micron sections of hippocampus, entorhinal cortex, midfrontal cortex, midtemporal cortex and inferior parietal cortex. The diagnosis follows the recommendation of the National Institute on Aging-Reagan criteria ^53^. Briefly, based on the scores of Braak stage for severity of neurofibrillary tangles and CERAD estimate for burden of neuritic plaques, a pathologic AD diagnosis requires an intermediate likelihood AD (i.e., at least Braak stage 3 or 4 and CERAD moderate plaques) or a high likelihood AD (i.e., at least Braak stage 5 or 6 and CERAD frequent plaques). **β-amyloid and tau pathology burden.** Quantification and estimation of the burden of parenchymal deposition of β-amyloid and the density of abnormally phosphorylated tau-positive neurofibrillary tangles levels present in the cortex at death (which we refer to as β-amyloid and tau pathology, respectively), tissue was dissected from eight regions of the brain: the hippocampus, entorhinal cortex, anterior cingulate cortex, midfrontal cortex, superior frontal cortex, inferior temporal cortex, angular gyrus, and calcarine cortex. 20µm sections from each region were stained with antibodies to the β-amyloid beta protein and the tau protein, and quantified with image analysis and stereology.

### Nucleus isolation and single nucleus RNA library preparation

Dorsolateral Prefrontal Cortex (DLFPC) tissue specimens were received frozen from the Rush Alzheimer’s Disease Center. We observed variability in the morphology of these tissue specimens with differing amounts of gray and white matter and presence of attached meninges. Working on ice throughout, we carefully dissected to remove white matter and meninges, when present. We expect this will help to reduce variability between tissue specimens. Working on ice throughout, about 50-100mg of gray matter tissue was transferred into the dounce homogenizer (Sigma Cat No: D8938) with 2mL of NP40 Lysis Buffer [0.1% NP40, 10mM Tris, 146mM NaCl, 1mM CaCl_2_, 21mM MgCl_2_, 40U/mL of RNAse inhibitor (Takara: 2313B)]. Tissue was gently dounced while on ice 25 times with Pestle A followed by 25 times with Pestle B, then transferred to a 15mL conical tube. 3mL of PBS + 0.01% BSA (NEB B9000S) and 40U/mL of RNAse inhibitor were added for a final volume of 5mL and then immediately centrifuged with a swing bucket rotor at 500g for 5 mins at 4°C. Samples were processed 2 at a time, the supernatant was removed, and the pellets were set on ice to rest while processing the remaining tissues to complete a batch of 8 samples. The nuclei pellets were then resuspended in 500ml of PBS + 0.01% BSA and 40U/mL of RNAse inhibitor. Nuclei were filtered through 20um pre-separation filters (Miltenyi: 130-101-812) and counted using the Nexcelom Cellometer Vision and a 2.5ug/ul DAPI stain at 1:1 dilution with cellometer cell counting chamber (Nexcelom CHT4-SD100-002). 20,000 nuclei in around 15-30ul volume were run on the 10X Single Cell RNA-Seq Platform using the Chromium Single Cell 3’ Reagent Kits v2. Libraries were made following the manufacturer’s protocol, briefly, single nuclei were partitioned into nanoliter scale Gel Bead-In-EMulsion (GEMs) in the Chromium controller instrument where cDNA share a common 10X barcode from the bead. Amplified cDNA is measured by Qubit HS DNA assay (Thermo Fisher Scientific: Q32851) and quality assessed by BioAnalyzer (Agilent: 5067-4626). This WTA (whole transcriptome amplified) material was diluted to <8ng/ml and processed through v2 library construction, and resulting libraries were quantified again by Qubit and BioAnalzyer. Libraries from 4 channels were pooled and sequenced on 1 lane of Illumina HiSeqX by The Broad Institute’s Genomics Platform, for a target coverage of around 1 million reads per channel.

### Pre-processing of snRNA-seq data

De-multiplexing, alignment to the hg38 transcriptome and unique molecular identifier (UMI)-collapsing were performed using the Cellranger toolkit (version 2.1.1, chemistry V2, 10X Genomics, for chemistry Single Cell 3’), and run using cloud computing on the Terra platform (https://Terra.bio). Since nuclear RNA includes roughly equal proportions of intronic and exonic reads, we built and aligned reads to a genome reference with pre-mRNA annotations, which account for both exons and introns. To remove the technical artifacts of ambient RNA molecules within the RNA profiles of each cell, we used the CellBender package. Briefly, CellBender builds an ambient RNA model by learning the background distribution of RNA in empty droplets, distinguishing cell-containing droplets from empty ones, and applying the model to achieve a background-free gene expression profile per cell (remove-background function, with 300 epochs and other default parameters). For every nucleus, we quantified the number of genes for which at least one read was mapped, and then excluded all nuclei with fewer than 400 detected genes. Genes with less than 15 reads mapped across all the nuclei were excluded. Expression values *E_i_*_,*j*_ for gene *i* in cell *j* were calculated by dividing UMI counts for gene *i* by the sum of the UMI counts in nucleus *j*, to normalize for differences in coverage, and then multiplying by 10,000 to create TPM-like values, and finally computing log_2_(TP10K + 1) (using the *NormalizeData* function from the *Seurat*^22^ package version 4). Next, we selected variable genes (using the *FindVariableFeatures* function in Seurat, setting the selection method to *vst*) and scaled the data matrix (using the ScaleData function from Seurat 3), yielding the relative expression of each variable gene by scaling and centering. The scaled data matrix was then used for dimensionality reduction and clustering.

### Dimensionality reduction and clustering

We used the scaled expression matrix restricted to the variable genes for Principal Component Analysis (PCA), using *RunPCA* method in Seurat (a wrapper for the irlba function), computing the top 50 PCs. After PCA, significant principal components (PCs) were identified using the elbow method, plotting the distribution of standard deviation of each PC (*ElbowPlot* in Seurat), choosing 30 PCs for analysis of all cells as for astrocytes and excitatory neurons, 25 PCs for inhibitory neurons and 20 for microglia, endothelial cells and oligodendrocytes. Scores from only these top PCs were used as the input to downstream clustering or visualization. The scaled data was embedded in a two-dimensional space using Uniform Manifold Approximation and Projection (UMAP) ^54^, based on the first significant PCs (as listed above). Clustering was performed using a graph-based approach: a *k*-nearest neighbor (*k*-NN) graph over the cells was calculated (using the *FindNearestNeighbors* function), followed by the Louvain community detection algorithm ^54,55^, which decomposes an input graph into communities to find transcriptionally similar clusters of cells (using the *FindClusters* function). The resolution of the clustering was selected using both cell-type markers and visualization of the UMAP embedding (using resolutions ranging between 0.1 to 0.6). In addition, we removed a group of nuclei that exhibited a neuronal signature that we suspected to be technically altered (expressed a low number of genes compared to other neurons, with missing expression of basic neuronal markers, and had a high signature of cytoplasmic specific genes). Cell populations were manually matched to cell types based on the expression of known marker genes as previously done^14,56^.

### Cell filtering and testing of potential technical artifacts

#### Removal of nuclei with high content of cytoplasmic-RNA and low nuclear-RNA

We manually detected clusters with high content of cytoplasmic RNA. The cytoplasmic RNA signature was defined as the top 400 differentially expressed genes between nuclei content and cellular content ^53^. Clusters that showed a decrease in nuclei content and an increase in cytoplasmic content were removed.

#### Doublet cells removal

For doublet detection and elimination, we clustered our data at high resolution, to generate multiple small clusters, and removed clusters enriched with suspected doublets. We used a combined approach of manual and automatic detection. First, we ran DoubletFinder ^57^ to identify high confident doublets that were removed from the dataset. Second, we identified *doublet clusters* defined as clusters with over 70% of nuclei classified as doublets, and we then excluded all nuclei within these clusters from the downstream analysis as well. Before removal, we validated that these clusters are *doublet clusters* based on a manual inspection of expression patterns of cell type marker genes, validating these clusters co-express markers of at least two different cell types. Of note, while at the cluster level, the automatic doublet analysis could have been done by expression of cell type markers, the automatic doublet analysis aided the identification of the doublet cluster and moreover aided the identification of doublet cells outside of doublet clusters. In the downstream analysis of sub-clustering of specific cell types, a second manual inspection for doublets was performed for expression patterns of cell type marker genes, excluding additional sub-clusters of doublet cells. Of note, some of these suspected doublet cells might have been cells with high content of ambient RNA, and were removed for technical reasons.

#### Testing technical artifacts driven by Batch or RIN and sex diversity

To rule out the possibility that the resulting clusters were driven by one or more covariates (e.g. technical effects of batch, and RIN or sex differences), we examined the distribution of samples within each cluster and the distribution of the number of genes detected across clusters (as a measure of nucleus quality). Additional filtration of low-quality cells and clusters was done following this initial clustering analysis, to remove low quality neuronal cells that could not be filtered out initially based on the number of genes detected, due to the cellular heterogeneity of the tissue, that includes both high RNA content cells (such as neurons) and low RNA content cells (such as microglia cells). To further validate that the clustering analysis is not driven by one or more covariates, we compared the sub-cluster assignment of all nuclei within the astrocytes, microglia and endothelial cell types (as in the described pipeline) to an alternative sub-cluster assignment after regression of each covariate in the scaled expression matrix. More specifically, for each cell type separately (astrocytes, microglia and endothelial cells) we first normalized the data as in the described pipeline, then regressed the effect of each covariate, independently, during the scaling of the expression matrix (using the *ScaleData* function in Seurat, with the vars.to.regress parameter). The downstream analysis of PCA and community detection clustering was performed as in the described pipeline. The cluster resolution was selected to match the number of clusters in the standard analysis. The two sets of cluster assignments were compared by calculating the overlap between each pair of clusters using the Jaccard index, showing the high overlap and the one-to-one match between the two sets of clusters.

### Sub-clustering and annotation analysis of glia, endothelial and neuronal cells

For each cell type that had a sufficient number of cells we performed sub-clustering analysis to reveal additional diversity of cell subtypes and cell states: astrocytes, microglia, endothelial cells, GABAergic inhibitory neurons and glutamatergic excitatory neurons (oligodendrocytes were analyzed by topic modeling, see the section *Topic modeling for oligodendrocyte cells*). To this end, the pipeline described for all cells was applied separately to each cell type including: normalizing the count matrix, finding variable genes, scaling the normalized count matrix, dimensionality reduction by PCA, graph clustering and dimensionality reduction by UMAP for visualization in 2D, and filtration of low-quality cells. For microglia, astrocytes and endothelial cells, initial annotations of subsets were done by known marker genes ^2, 4–6,14^. Microglia subsets were further annotated by comparison to a dataset of live microglial cells^6^ and all subsets were compared to existing lower resolution clustering (see the section *Comparison to published datasets of snRNA-Seq and scRNA-Seq*).

To annotate neuronal subtypes and predict the cortical layer of each neuron, we performed an automatic annotation by running the SingleR package against a fully annotated and detailed cortical single nuclei RNA-seq dataset from the Allen Brain Atlas^11,52^. The annotation was performed separately for the inhibitory and excitatory neurons (based on an initial annotation by known marker genes^11,56^) and predicted both the neuronal layer level and the neuronal subtype. We used the “fine tuned” label within the Allen Atlas in order to determine the best cortical layer fit. Annotating neuronal subtypes was first done by known marker genes, assigning neuronal cells to excitatory and inhibitory classes. The learned regression model was then applied to our neuronal snRNA-seq data, inferring for each nuclei the best label of cell subtype and cortical layer. Annotations in our dataset were done on the cluster level, based on the highest scoring label across all cells. The automatic annotations were validated by the expression of known marker genes^11,56^.

### Topic modeling for oligodendrocyte cells

We modeled the cell state diversity in oligodendrocyte cells by topic modeling^54^, using Latent Dirichlet Allocation (LDA). Topic modeling was performed on the normalized data matrix, reduced to the oligodendrocytes variable genes. To assign topics and score cells for each topic, we used the CountClust package in R, which calculated the grade of membership for four topics (using the GoM function, which was run on the scaled expression matrix with tolerance 0.01). To find the optimal number of topics and tolerance levels, we ran a grid of parameters of topic numbers and tolerance levels, and visualized topic scores on the UMAP. The results were robust to the choice of parameters, yet we favored a small number of topics given the number of cells and samples in our data. A higher number of topics often captured cells of one individual and would thus require more individuals for reproducibility. To select the genes highly associated with each topic, we used the Kullback-Leibler divergence (KL), which measures the difference between two probability distributions, applied to the distribution of the topic weights and the distribution of gene expression level across cells (using the ExtractTopFeatures function with default parameters). As we were interested in positive association between genes and topics, we filtered out genes with negative correlation to the related topics, for downstream analysis such as validations and pathway enrichment.

### Densities plots to visualize distribution of cellular populations and gene expression in 2D embeddings

For clear visualization of the distribution of cell populations or distribution of the expression levels of a gene, in dense 2D graphs of multiple cells, we used a Gaussian kernel approach to plot the density of cells of a specific subset, a score per cell, or a gene expression level. We used this approach to plot the expression of marker genes and to plot scores of the four topics of oligodendrocytes in Fig.3. For discrete traits, we scored each cell by calculating the density of neighbors sharing the same trait, calculating the Gaussian kernel adjacencies of the 2D UMAP embedding for all other cells (using the Gauss kernel from the ‘KLRS’ package in R, with sigma = 1.5). We filtered out cells with an adjacency measure > 0.0005. For the remaining cells, we calculate the proportion of cells assigned to individuals with the relevant phenotype, weighing them by their Gaussian distance to the cell. Finally, we normalize the obtained measure of the cell by the total distances to the remaining cells. In the continuous case, we use a similar approach, considering all neighboring cells weighted by their individual expression level.

### Differential expression and pathway analysis

Differentially expressed signatures were calculated using a GLM-framework (*FindAllMarkers* function, test.use = “MAST”) and controlled false-discovery rates (FDRs) using the Benjamini–Hochberg procedure, to find genes that are down or upregulated within each cluster compared with the rest of the nuclei in the dataset including genes with less than 5% FDR. Genes were required to be expressed in at least 10% of nuclei in the given cluster, and to have at least 0.25-fold lower average expression in all other cells outside of the cluster. To calculate differentially expressed genes in oligodendrocytes, we used nuclei that scored more than 0.5 to one of the topics and addressed it as a regular hard assignment. The differential expression signatures were tested for enriched pathways and gene sets (function *compareCluster* in the clusterProfiler package in R), and corrected for multiple hypotheses by qvalue. Results with qvalue < 0.05 were reported as significantly enriched pathways. Gene sets and pathways were taken from the KEGG and Gene Ontology (GO) resources^58^.

#### Shared pathways analysis

To detect shared enriched biological pathways among a group of subsets, we used the pathways calculated separately for each subset, and selected pathways that were enriched in at least 3 cell subsets. To overcome the redundancy within the enriched pathways, the shared pathways were clustered by their gene similarity to capture discrete annotations that represent a non-redundant set of enriched pathways. More specifically, for each enriched pathway we chose the genes *within* the pathway that are also differentially in one or more of the relevant cell subsets. We then clustered the pathways by hierarchical clustering (pheatmap function) using the pairwise Pearson Correlation distance computed over the gene space. Each pathway cluster was named manually according to the pathways within the cluster.

### Comparison to published datasets of snRNA-Seq and scRNA-Seq

We mapped our microglia snRNA-Seq to the recently published dataset of scRNAseq of live microglia cells from fresh autopsy and surgically resected human brain tissue ^6^. The mapping was performed using Canonical Correlation Analysis in Seurat with standard parameters (3,000 genes and 20 canonical components) to align the snRNA-Seq and scRNA-seq data. We then used a naive Bayes classifier trained on the scRNA-seq-derived live microglia clusters in canonical correlation component space. This classifier was used to predict the class membership of the CCA-transformed snRNA-Seq nuclei profiles, and each nucleus profile was assigned to the single-cell live microglia cluster with the maximum prediction value. Applying the same CCA-based approach we mapped our cell type clusters and all of our cell sub-clusters to four recently published datasets of snRNA-seq of human brains^2,3,5,10^.

### Estimating major cell types, cell subsets and topics proportions from bulk data by CelMod

We developed a regression-based consensus model (CelMod) to extend our snRNA-seq-derived cell class and subset estimates to bulk data, leveraging matched bulk and snRNA-seq data from the same 24 donors.

We train the regression model as follows: (**1**) Filter genes to include only those that have at least 100 counts across all nuclei of the cell type of interest, and a mean counts per million value > 10, (**2**) Perform a linear regression on each gene separately for each cell cluster of interest, using its expression as the dependent variable and the proportion of that cluster in each snRNA-Seq sample in the training set as the independent variable; (**3**) For each gene, use the regression model to calculate the predicted proportion of each cell type, normalizing their sum to 1; (**4**) Rank genes by the 90^th^ percentile of the absolute value of the error between predicted and training proportions, for each cell type; and (**5**) Select the number of top-ranked genes (constant for each cell cluster) to use for deconvolving a new bulk RNA-Seq sample; this number of genes, the only tunable parameter, is selected based on cross-validation, as described below.

To determine the optimal number of genes to use for the prediction, we use 5-fold cross-validation using 80% of the data for training and 20% as the validation set. The validation sets are mutually exclusive, such that after 5 runs, the proportions in every bulk sample have been “predicted” once. This cross-validation is run using 3 to 100 ranked genes (from step 3 above), and the optimal gene number is selected as that which minimizes the mean of the 90^th^ percentile errors for each cell group in all samples. This gene number is then used for deconvolution predictions in the larger bulk RNA-seq data set, for a given set of cell clusters.

We run the algorithm iteratively, starting at the “top level”, with the broad cell classes (glutamatergic neurons, GABAergic neurons, astrocytes, oligodendrocytes, OPCs, microglia, endothelial cells, and pericyte), and then again for the subtypes within each of the cell classes. For the broad cell classes, the proportions are based on the total nuclei per sample. For the subtypes, the proportions are normalized to the total nuclei from the broad class of interest - for example, for astrocyte subsets, the proportions for the training (and thus the prediction) are normalized to the total number of astrocyte nuclei per sample. This allows us to directly model both the overall and subtype-level composition of the bulk tissue, especially for cell types that comprise a small fraction of the overall population (such as the endothelial or microglial subtypes). For the oligodendrocyte signatures, which are modeled as topics instead of discrete clusters, we sum the weights for each given signature over all nuclei from a given sample, and then normalize these sums so that they add up to 1 for a given sample. This reflects a “proportion of topic weights” per sample, as opposed to a strict proportion that can be calculated for the discrete cell types. Finally, for microglia, we ensured the model training was robust, by only including donors with at least 25 total microglia.

The performance of this algorithm on the validation set (20% of the data, with 5-fold cross-validation) is shown in **Fig. 4c-e and Extended Data Fig. 6a-c**, as well as the correlation structure between cell subsets at **Fig. 5d,e and Fig. 6c**.

We applied the inferred CelMod model to a set of 106 bulk RNA-seq profiles from a different independent cohort from the Mount Sinai Brain Bank (MSBB^32^). The samples were pre-processed similarly to the ROSMAP dataset (described in ^48^), then we applied CelMod model learned on our snRNA-seq data to estimate the major cell types, cell subsets and topics proportions in each bulk sample. The cell subset estimates from this independent MSBB cohort were used to validate the associations of cell subsets proprtions to AD-associated traits (**Extended Data Fig. 6f**) and the correlation structure of cellular subsets (**Extended Data Fig. 7a**). The analysis followed the same methods as described for the bulk RNA dataset of the ROSMAP cohort with one exception. Due to constraints of the available metadata we used in the MSBB cohort the CDR measurement that quantifies the level of cognitive decline, and the BRAAK score that quantifies the level of tangles burden (instead of the cognitive decline rate and the tangles load used for the ROSMAP cohort, respectively).

We compared the performance of CelMod to 3 bulk deconvolution methods: DCQ, DeconRNAseq, and dtangle, with standard parameters for each method. Each of these methods takes bulk RNA-seq and uses reference profiles of the snRNA-seq-derived clusters to estimate proportions. We ran each of these methods both for “top level” cell classes, as well as with the full-resolution subclusters, and calculated Spearman correlation r-values between the predicted proportions and the actual snRNA-seq-derived proportions. We note that none of these comparator deconvolution methods were designed specifically to recapitulate snRNA-seq proportions, and so do not take into account cell type-specific biases due to dissociation or other aspects of the protocol.

### Statistical analysis to assign cell types, subtypes, cell states and topic models to AD related traits

We analyzed five major pathological and cognition hallmarks of AD, collected as part of the ROS/MAP cohort, including: primary neuropathological and cognitive phenotypes. The two clinical traits were a clinical diagnosis of AD dementia proximate to death (clinical AD) and a continuous measure of cognitive decline over time quantified as a per-subject slope of the cognitive decline trajectory from a linear mixed-effects model ^50^. The three pathology variables include continuous measures of tau tangle pathology density and β-amyloid burden (both averaged over multiple regions) and a binary diagnosis of pathologic AD (as previously described^59^). Details on the traits can be found in the section: *Experimental design: AD traits in the ROS/MAP cohorts*.

To test the statistical associations between neuropathological phenotypes and cell types or cell subsets proportions, we performed multivariable linear regressions, modeling cellular proportions as the outcome, and neuropathology as the independent variable. In each analysis, we adjusted for age, sex, and RIN score as covariates to remove potential confounding, and corrected for multiple testing, by calculating the false discovery rate (FDR) for each set of pathology analyses. The analysis was done for each of the five traits separately (amyloid, tau, cognitive decline, pathologically determined Alzheimer’s disease, and post mortem clinical diagnosis of Alzheimer’s disease), across all cell types (astrocytes, oligodendrocytes, endothelial, inhibitory neurons (GABAergic), excitatory neurons (glutamatergic), microglia, pericyte, and oligodendrocyte progenitor cells (OPCs)), or across all cell subsets (5 astrocytes, 6 endothelial subsets, 7 GABAergic neurons, 10 glutamatergic neurons, 5 microglia subsets, and 4 oligodendrocyte topics).

### Cellular Communities

To find co-occurring cellular populations across individuals, we generated a network of cellular communities, defined as a set of cell populations (cell types, subtypes, cell states or expression programs) that have coordinated variation of proportions across individuals (for topics we use the proportional weights instead of the subset’s proportion). We applied this approach to define cellular communities from our snRNA-seq dataset (n=24 individuals) and from the CelMod estimated proportions (n=638 individuals), across 6 broad cell classes (astrocytes, microglia, endothelial cells, oligodendrocytes, inhibitory (GABaergic) and excitatory (glutamatergic) neurons) for which we had a defined sub-clustering (splitting each cell type to subsets capturing distinct cell subtypes or cell states), or defined topic models (splitting each cell type to distinct expression programs, without the need for a discrete assignment of cells to clusters). When applying this approach to the estimated cellular proportions by the CelMod algorithms in 638 individuals, we included the estimations for the oligodendrocyte cells, which allowed for a distribution of weights across the topics for each cell. To detect cellular communities, we followed these four steps: **(1) Cellular proportions.** Given a classification of cells (in the current analysis the subsets are cell subtypes for neurons, cell states for glial and endothelial cells, and expression programs (topic models) for oligodendrocytes), we calculated for each cell class and for each individual: the frequency of all cell subsets *within* each cells class (i.e. the proportion of cells within a subset out of the total number of cells/nuclei in that broad cell class or sum of all topic weights, for oligodendrocytes). Next, we appended the cell subset proportions across *all* cell classes into a *combined frequency matrix*. Of note, for the 638 individuals we used the estimated proportions by CelMod. **(2) Correlation matrix.** We calculated the pairwise Spearman correlation coefficient over the frequency of each cell subset across individuals, created a pairwise correlation matrix and clustered it by hierarchical clustering (using the pheatmap function in R, based on the distance metric of 1-Pearson correlation). **(3) Cellular network.** We transformed the correlation matrix into a graph, where each cell subset is a node, and edges connected every two nodes if their absolute pairwise Spearman correlation value was at least 0.4 and p.value < 0.05. Each edge was assigned a positive or negative sign. In order to maintain a consistent common layout between the networks derived from 24 and 638 individuals, the graph was manually laid out, using the igraph package (in R), and negative edges were added for visualization (setting the parameter layout = layout_with_fr). **(4) Associating cellular communities to AD-traits.** Each cellular subset (nodes in the network) was first associated to AD-traits by multivariable linear regression (see details in the section *Statistical analysis to assign cell types, subtypes, cell states and topic models to AD related traits*), associations with FDR<0.01 were considered significant and assigned as positive (beta>0) or negative (beta<0) associations, while association with FDR > 0.01 were considered as neutral. Next, **significant connected components associated with AD-traits were termed communities.** To test the significance of the association: For each AD trait (cognitive decline, tangles burden or amyloid burden), we calculated the connectivity score within three sets of subsets (nodes in the network): positively associated, negatively associated or neutral to this trait, and between these sets. We defined the connectivity score for a set of nodes in the network to be: the differences between the numbers of positive and negative edges divided by the total number of possible edges between them (edges as defined in the graph). To assess the statistical significance of high and low scores, we applied a permutation test on the subset labels within the network. The empirical p-value is the proportions of permutations (out of 10000) that lead to a higher/lower score.

Of note, since we used only 4 topics that are largely disjoint sets of cells for the oligodendrocytes, a hard assignment for an oligodendrocyte to a specific topic (by the maximum weight, thus transforming the data to discrete clusters), did not substantially impact the results compared to the continuous topic weight. However, the framework proposed here applies more generally to any level of topic modeling or other soft assignment of cells to continuous cell states and expression programs.

#### Testing potential layer bias underlying the network analysis

To exclude the possibility that correlations between subsets are induced by cortical layer proportions driven by a dissection bias of the cortical samples, we compared the correlation matrices of two distinct groups. Excitatory neuronal subsets largely formed two opposing independent connected components in the graph, one consisting of upper cortical layer neuronal subsets (Exc.1, Exc.2, Exc.3, layers 1-4) and the other of lower layer neuronal subsets (layers 4-6). Given the intrinsic association between pyramidal neuron subsets and cortical layers we cannot completely exclude the possibility that this partition is driven by a dissection bias of the cortical samples, yet, we can do that for other cell subsets (inhibitory, endothelial and glial subsets). We first divided individuals based on their CelMod estimated proportion of Exc.1, which divided the 638 individuals into 2 distinct groups: High levels of Exc.1 (> 0.5) or low levels of Exc.1 (<= 0.5). Next, we calculated the pairwise Spearman correlation coefficient matrices of the CelMod proportions of all cell subsets across the two sets of individuals separately: low levels of Exc.1 (n=371 individuals) and high levels of Exc.1 (n=267 individuals). Next, we calculated the differences of the two pairwise correlations matrices, showing the partition mainly affects excitatory neurons, for which the absolute change was higher than 0.2.

#### Comparison of the snRNA-Seq network and the CelMod network

We compared the snRNA-Seq network (24 individuals) and the CelMod estimated cellular network (638 individuals, as in **Supplementary Fig. 5b,c**). First, we calculated the statistical significance of the similarity between the snRNA-Seq network and the CelMod network by an empirical p-value: We performed 10,000 random permutations of the combined frequency matrix over all cell subsets in the 638 individuals (estimated by CelMod), shuffling the values within each cell cluster, such that the frequency of cell subsets will sum to 1 for each cell type. We calculated a correlation matrix for every permutation, and a similarity score between the snRNA-seq correlation matrix and the permutation matrix using Jennrich’s score^60^ (cortest.jennrich function in R). An empirical p-value (p<0.001) was assigned to the original CelMod estimated proportions, since the non-permutation matrix consistently got the lowest score compared to the permuted-matrix.

### Analysis of proteomic data

Shotgun bulk proteomic data of 400 individuals from 400 aging participants of the ROSMAP cohort (recent publication^35,36^), with 160 individuals overlapping the 638 individuals profiled by bulk RNA-seq used in our study. The protein expressions were normalized by the median of the proteins expression per sample. We used the log2 transformed levels (as described in ^37^) of protein markers of cell subsets of interest we validated the following: (**1**) Estimated proportions from bulk RNA-seq of cell subsets by CelMod; (**2**) Associations between cell subset proportions to AD-associated traits; (**3**) Correlation structure between propotyiond of different cell subsets. In all these analyses, we first chose differential marker genes based on the RNA expression in the snRNA-seq data, as described for the differentially expressed genes, but requiring in addition a low or no expression outside of the cell population of interest, in and outside the same cell type. Next, we filtered out marker genes that were not detected or were lowly expressed on the protein level in the proteomic dataset. Finally, cell subsets that didn’t have sufficient protein markers expressed were not considered for the validation analysis. Of note, the basic protein and RNA levels of multiple genes did not correlate well which limited this analysis.

### Immunohistochemistry and Spatial transcriptomics

Formalin-fixed *post-mortem* brain tissues were obtained from Rush University Medical Center. As part of these studies, all participants consent to brain donation at the time of death. 6µm sections of formalin-fixed paraffin-embedded (FFPE) tissue from the cortex frontal were stained with NEF (Sigma, N2912). Heated-induced epitope retrieval was performed using citrate (pH=6) using microwave (800W, 30% power setting) for 25 min. The sections were blocked with blocking medium (3% BSA) for 30 min at Room Temperature, then incubated with primary antibody anti-NEF prepared in 1% BSA for overnight at 4°C. Sections were washed three times with PBS and incubated with fluochrome conjugated secondary antibodies (Thermo Fisher) for one hour at RT. Anti-fading reagent with Dapi (P36931, Life technology) was used for coverslipping. For each subject, 30 images of cortical grey matter at magnification x20 (Zeiss Axio Observer.Z1 fluorescence microscope) were taken in a zigzag sequence along the cortical ribbon to ensure that all cortical layers are represented in the quantification in an unbiased manner. The acquired images were analyzed using CellProfiler and Cellprofiler Analyst developed by Broad Institute. We estimated the proportions of a broad cell class or signature (neurons, microglia and GFAP+ astrocytes) from the images as the fraction of nuclei stained with the marker of interest (NeuN, IBA1, or GFAP, respectively for neurons, microglia, and GFAP+ astrocytes) out of all nuclei stained by DAPI.

For spatial transcriptomic, the cerebral cortex and the underlying white matter of fresh frozen brain tissue was obtained from New York Brain Bank for six aging individuals (NYBB) and dissected on dry ice. The samples from each subject was prepared into 10mmx10mm size OCT and stored in −80°C until the experiment. Samples were sectioned at 10 µm of thickness in duplicate onto a slide containing capture probes. Each tissue section contained all six cortical layers. Sections were fixed with cold 100% methanol for 30 min and then stained with H&E for 7 min at Room Temperature. Sections were scanned using a Leica Microscope. After image capture, tissue sections are permeabilized to induce cDNA synthesis and a cDNA library is generated where each molecule is tagged with its spatial barcode. The permeabilization time was optimized for the ST Visium platform and ensured that the RNA quality in fresh frozen brain tissue was sufficient for ST data generation. Only tissue sections with a RIN score above 7 were selected. The libraries were then sequenced and barcoded cDNA libraries were aligned with H&E image of the tissue section using the Space Ranger software. The quantification of each gene for each spot was analyzed using the Seurat Package V.4.1.0. Visualization of spatial distribution of genes of interest across the six tissue sections were computed using the SpatialFeaturePlot function from Seurat V.4.1.0.

### Ligand-receptor analysis

We searched for ligand-receptor expression as an indication for interactions between different cell subsets, within cellular communities and between cellular communities. Focusing on all cell subsets within the *cognitive decline* community (Inh.1, Inh.7, Oli.4, Ast.4, End.2) and within the *cognitive non-impaired* community (Inh.3, Oli.1, End.4, End.1, Ast.2) and between the communities. Ligand and receptor interactions were assembled from published resources^35,36^ and manually curated. We searched for significantly expressed ligand-receptor pairs (LRP) between pairs of cell subsets using CellPhoneDB 2.0 ^37^. Briefly, this method computes the average expression of a ligand-receptor pair for each pair of cell subsets provided as a meta file. It then computes an empirical p-value to the specificity of this interaction by permuting cell subsets identity 1,000 times and comparing the real average expression against this null distribution.

Next, we found the significant interacting pairs of cell subsets, based on their overall number of LRPs. To calculate the statistical significance of the strength of estimated interaction between pairs of cell subsets, we detonated each cell-subset pair ligand-receptor interactions as edges in a directed cell-cell interaction graph. We performed a permutation test (10,000 permutations) to calculate an empirical p-value for the number of interactions found for a specific pair of cell subsets. The permutations maintained the total number of edges, by randomizing the ligand- and receptor-cell subset assignments.

To find the community specific differential LRPs, we defined a community specific LRPs as a ligand-receptor pair that were found to be both statistically significant expressed in the respective pair of cell subsets, but the ligand or the receptor were found to be differentially expressed within these subsets compared to all other cells of the same type (using the list of differentially expressed genes as defined in the section Differential expression and pathway analysis). A similar permutation strategy was used to compute an empirical p value for the strength of interactions to find significantly interacting pairs of cell subsets. A similar statistical approach and framework was used to define a strict list of community specific LRPs, in which both the ligand and the receptors were differentially expressed, and pairs of cell subsets that were enriched with such specific LRPs (**Extended Data Fig. 8a**).

### Causal modeling between cell subtype proportions and AD endophenotypes

To assess plausible causal relationships among CelMod-inferred cell subtype proportions and AD endophenotypes (Aβ, tau, and cognitive decline), we performed a series of linear regressions followed by mediation analysis. As purely data-driven causal inferences among cross-sectional traits is limited, we built our models on the known sequence of AD progression that starts with Aβ accumulation, that is followed by tau aggregation and cognitive decline. That is, we assumed that the direction of effect is from Aβ to tau, and that cognitive decline is a downstream consequence of tau accumulation, not the other way.

We focused on cell subtypes that are associated with tau pathology that were also nominally associated with Aβ (p<0.05). We tested the following linear model to assess the cell subtypes’ position in the causal chain including Aβ and tau, testing different scenarios for the effect of each cell subset proportion: upstream of Aβ, between Aβ and tau, or downstream of tau:

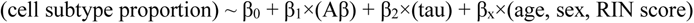

If β_1_ is attenuated after including tau in the model, tau is statistically either a confounder (independent effect on both Aβ and cell subtype proportion) or a mediator (relaying Aβ effect on cell subtypes)^61^. However, we assume that the direction of effect is from Aβ to tau based on the literature^39,40^, and thus tau will be inferred to be a mediator not a confounder, and cell subtype proportion will thus be inferred to be an outcome downstream of tau (the mediator). Since the first linear model indeed inferred the majority of subtype proportions to be downstream of tau, therefore, we set a following mediation model in this case:

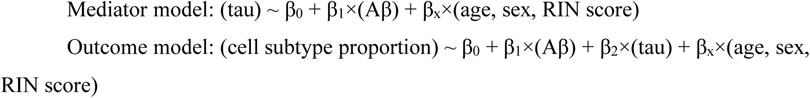

Here, Aβ is the independent causal variable, tau is the mediator, and cell subtype proportion is the continuous outcome (dependent variable). We used R “mediation” package^62^ to perform mediation analysis using this model. Mediated (indirect) effect, direct effect, and proportion mediated were estimated using non-parametric bootstrap method with 10,000 simulations.

Similarly, for the cell subtype proportions that are likely downstream of tau, we assessed their impact on the downstream cognitive decline. Here, we assumed that cognitive decline is downstream of brain cell subtype proportion changes, as this is a more plausible direction of causation (rather than a model assuming that poor cognitive performance leads to brain cell subtype changes). Here, the mediation model we used is:

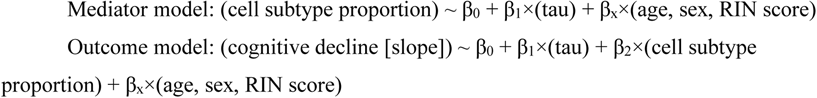

Here, tau is the independent causal variable, cell subtype proportion is the mediator, and cognitive decline (slope) is the continuous outcome (dependent variable). Mediation analysis was performed using the same package and setting as above.

Then we estimated the joint effect of cell subtype proportions that mediate tau effect on cognition comparing β_1_ (effect size of tau) of the following two linear models:

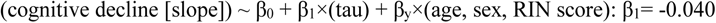

## DATA AND SOFTWARE AVAILABILITY

Raw and processed snRNA-seq data is deposited in the SYNAPSE database (https://www.synapse.org). Other ROSMAP data can be requested at the RADC Resource Sharing Hub at https://www.radc.rush.edu. Processed data, CelMod model for deconvolution and code will be available at the time of publication in the Single Cell Portal (https://singlecell.broadinstitute.org/single_cell) and in the GitHub.

## SUPPLEMENTARY TABLES

**Supplementary Table 1** - Samples details and phenotypes.

**Supplementary Table 2** - Cell annotations clustering and projids

**Supplementary Table 3** - Differentially expressed genes for cell subtypes/states.

**Supplementary Table 4** - CelMod estimated cellular proportions for 638 individuals

**Supplementary Table 5** - AD-traits associations to cell types and cell subsets proportions

## ACKNOWLEDGEMENTS

We thank the individuals who have generously donated their brain to research through the RUSH University Alzheimer’s Disease Center. We thank members of the Habib and DeJager labs, specifically, Adi Ravid and Or Gold; This work was supported by: grants from the National Institute of Aging (AMP-AD U01 AG046152, U01 AG061356, RF1 AG036042)(P.L.D.&D.A.B), the Chan Zuckerberg Initiative’s Neurodegeneration Challenge Network (CS-02018-191971)(P.L.D.), the Klarman Cell Observatory (A.R.), Alon Fellowship and Myers Foundation (N.H.), Israel Science Foundation (ISF) research grant no. 1709/19 (N.H.), The European Research Council grant no. 853409 (N.H.), MOST/IL grant no. 3-15687 (N.H.). F.Z. and A.R. are Investigators of the Howard Hughes Medical Institute. N.H is a Goren Khazzam senior lecturer in neuroscience. ROSMAP is supported by NIA grants P30AG10161, R01AG15819, R01AG17917, U01AG46152, and R01AG61356.

## AUTHOR CONTRIBUTIONS

P.L.D. and N.H. designed the study; C.M. prepared the single nucleus libraries and performed sequencing; A.C and V.M. performed the computational analysis, with guidance of N.H., V.M., P.L.D. and help from G.G.; I.H. performed the analysis of cellular interactions under the guidance of E.Y.L.; H.Y. and C.W. performed the statistical and modeling analysis under the guidance of P.L.D.; P.L.D., N.H., V.M., A.C. D.A.B., E.Y.L. and A.R. wrote the manuscript, and all co-authors edited it for critical comments. D.A.B. is PI of the parent ROS and MAP studies and obtained funding and performed study supervision. P.L.D., D.A.B and N.H. deposited data in Synapse and the RADC Resource Sharing Hub.

