## Extended Data Figures for "Multi-cellular communities are perturbed in the aging human brain and Alzheimer’s disease"

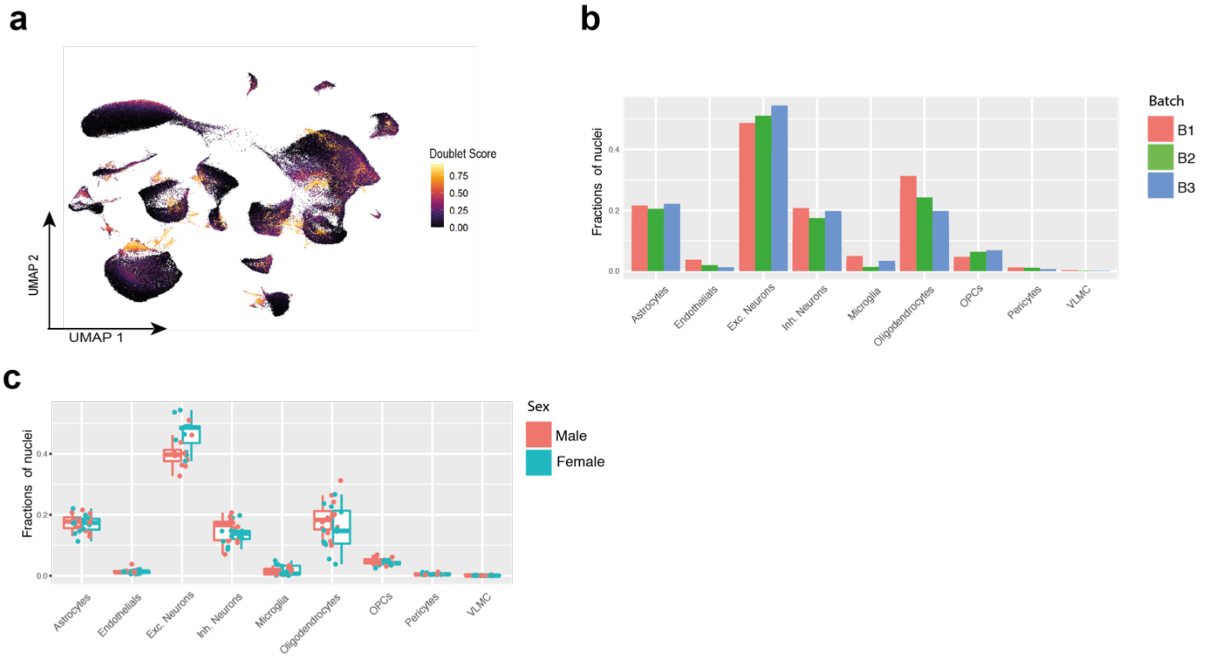


**Extended Data Figure 1. A cellular-molecular map of the human aging DLPFC: Quality controls (Accompanying Figure 1)** (**a**) High quality 172,659 nuclei libraries generated across 24 post-mortem samples of the DLPFC brain region of aging individuals. Nuclei (dots) colored by the doublet score attributed by DoubletFinder (see Methods). (**b**) Distribution of cell type frequencies across batches. Bar plot showing the fraction of nuclei per cell type for each batch. (**c**) Distribution of cell type frequencies across sex. Boxplot showing the fraction of nuclei per cell type for males (blue) and females(red). Box: 75% and 25% quantiles. Line: median. Dots: individual samples.

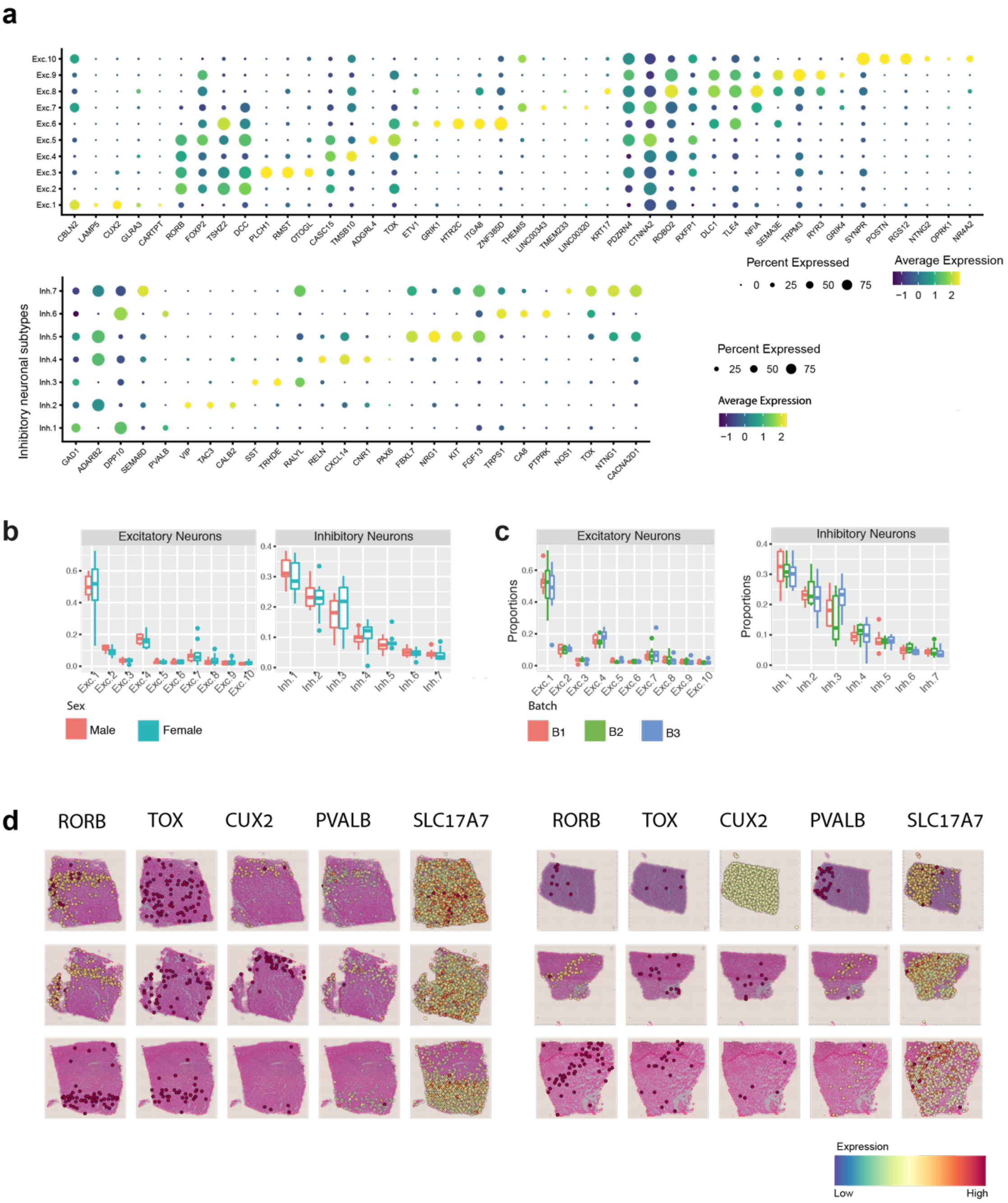


**Extended Data Figure 2. Quality control of neuronal subtypes and their cortical layer specificity (Accompanying Figure 2).** (**a**) Distinct expression of known and *de novo* marker genes in excitatory neuronal subtypes (top) and inhibitory neuronal subtypes (bottom) as assigned by our clustering analysis. Mean expression level in expressing cells (color) and percent of expressing cells (circle size) of selected markers in each neuronal subtype (rows) of marker genes. (**b-c**) Distribution of neuronal subtype frequencies across sex (b) and batches (c). Boxplot showing the fraction of nuclei per neuronal subtype for sex (b) or batch (c): Box, 75% and 25% quantiles. Line, median. Dots, individual samples. (**d**) Marker genes of neuronal subtypes exhibit a spatial organization at distinct layers within DLPFC slices. Spatial transcriptomics across 6 slices from 3 individuals for five marker genes (RORB, TOX, CUX2, PVALB, SLC17A7). Note variable orientation of slices. Complementary images to **Fig. 2c**.

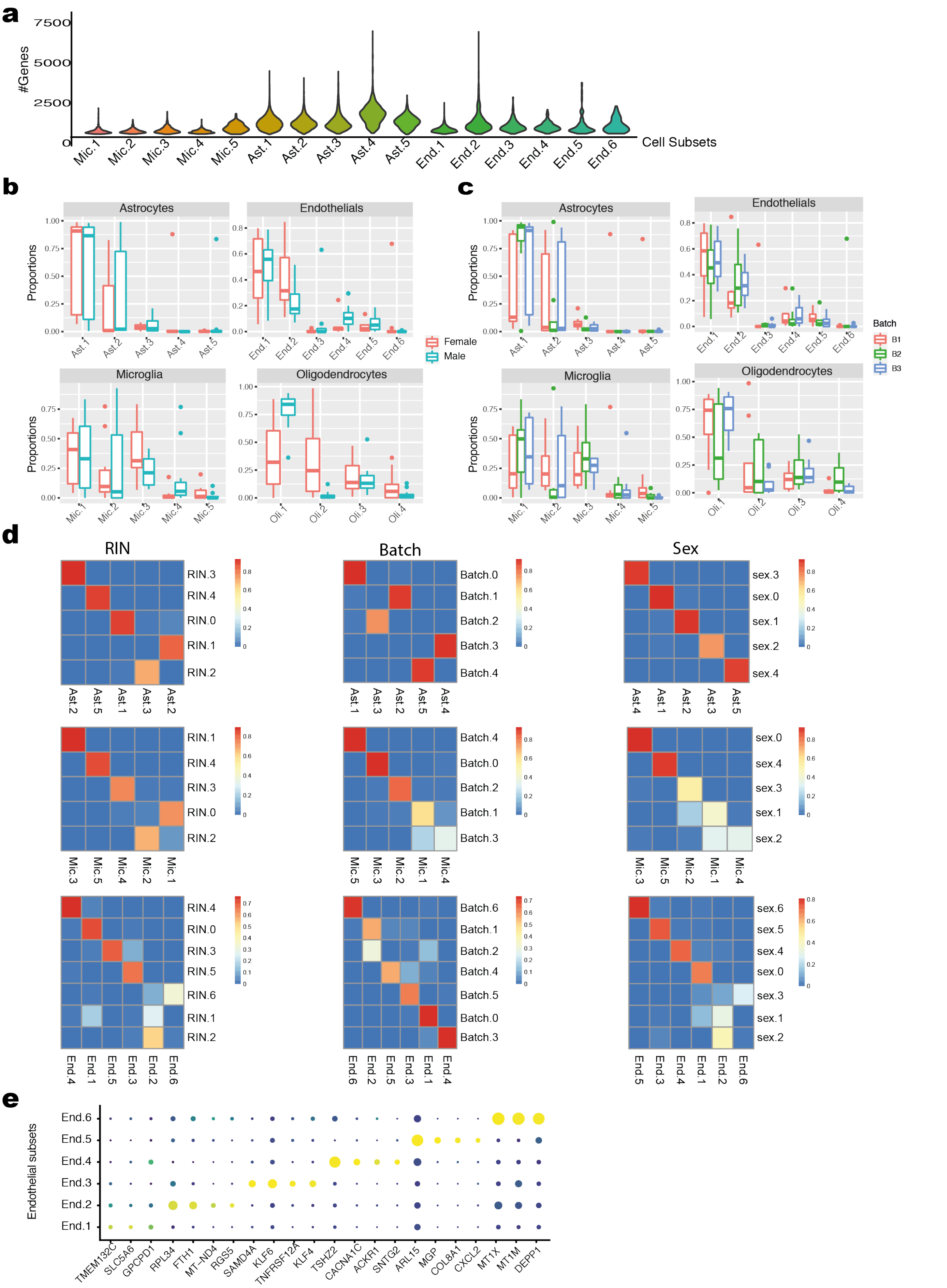


**Extended Data Figure 3. Quality control of glia subsets (Accompanying Fig. 3).** (**a**) Distribution of number of genes across microglia, astrocytes and endothelial clusters (denoted as subsets). (**b-c**) Distribution of non-neuronal subsets frequencies across sex and batches. Boxplot showing the fraction of nuclei per cell neuronal subtype for sex (b) or batch(c): Box, 75% and 25% quantiles. Line, median. Dots, individual samples. (**d**) RIN, sex and batch do not affect the sub-clustering of astrocytes, microglia and endothelial cells. Heatmaps of Jaccard score comparing overlaps of cells (color scales) between assignment of de-novo clusters after regression of the confounding variables from the expression matrix (rows) compared to the clusters in this study without such correction (columns) (**Methods**). **(e)** Endothelial subsets express unique markers. Dot plot of the mean expression level in expressing cells (color) and percent of expressing cells (circle size) of selected marker genes across endothelial subsets.

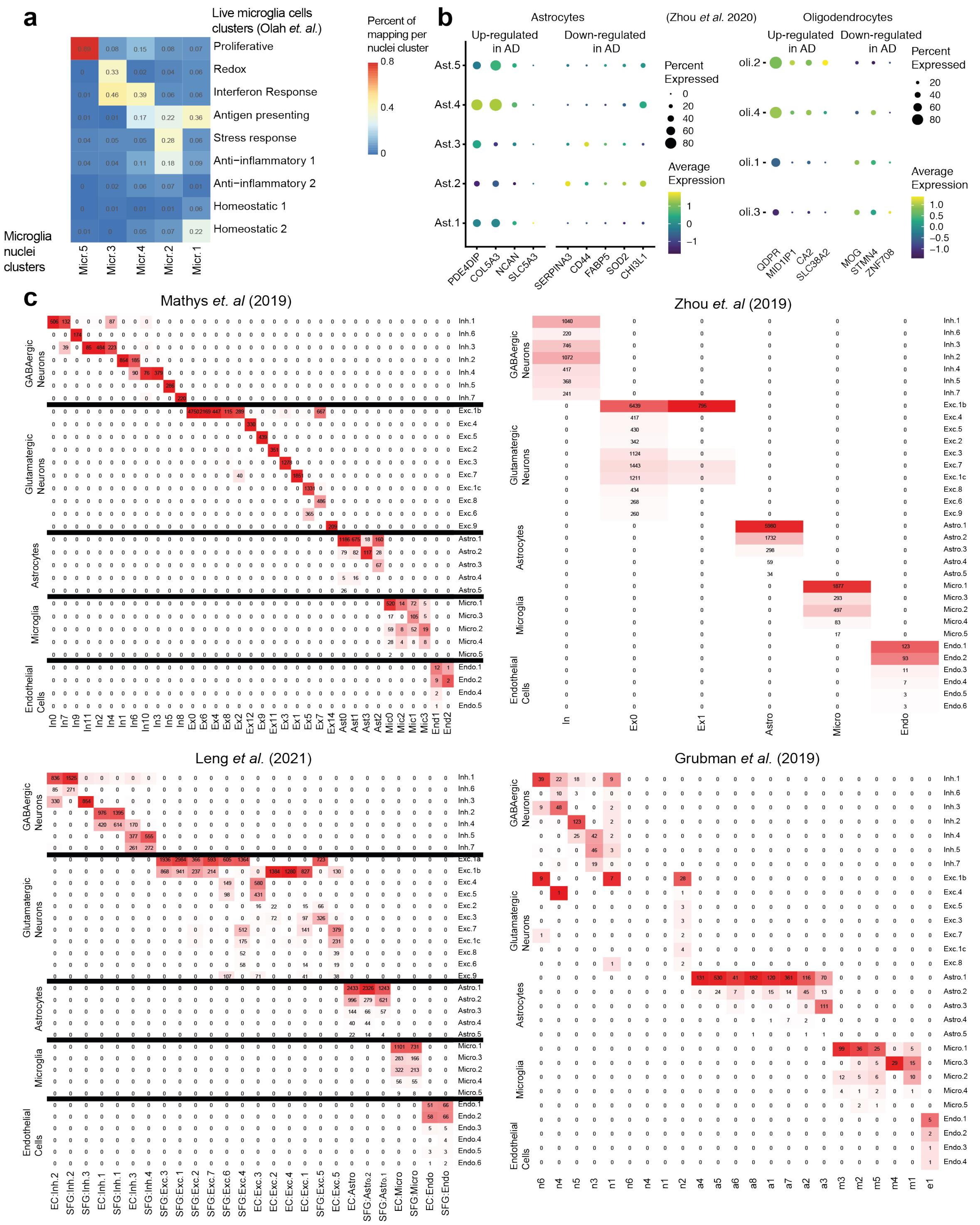


**Extended Data Figure 4. Comparison of cell clusters to previous studies (Accompanying Fig. 3).** (**a**) Clusters of microglia nuclei from snRNA-seq match published live microglia cell clusters from scRNA-seq. The proportions (color scale, scaled per column) of nuclei per cluster (columns) mapped to each scRNA-seq cell cluster according to the best prediction (rows, Methods). (**b**) Mean expression level in expressing cells (color) and percent of expressing cells (circle size) across cell subsets (rows) of previously described up-regulated and down-regulated genes in AD brains compared to healthy individuals, as defined by Zhou *et al*[2,3,5,10] for astrocytes (left) and oligodendrocytes (right) . (**c**) Nuclear-derived model is consistent with earlier, lower-resolution models across different cell types. Heatmaps (color scale) of assignment of nuclei from 24 individuals (rows) to published subsets (Methods) from 4 previous models of DLPFC.

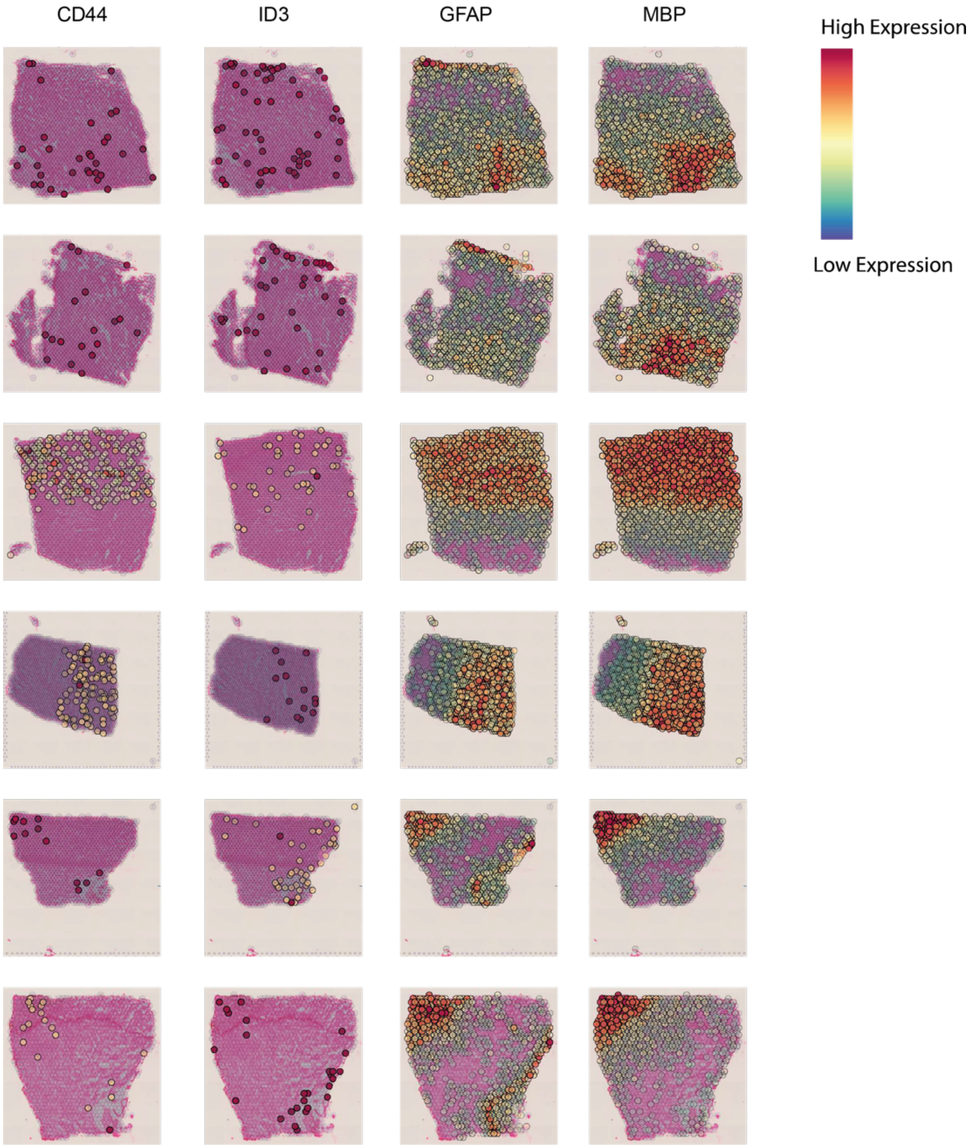


**Extended Data Figure 5. Spatial transcriptomics of glial markers (Accompanying Fig. 3).** Spatial transcriptomics of markers of glial cell types and states exhibit a spatial pattern across cortical layers matching DLPFC and white-matter anatomy. MBP (oligodendrocyte marker), GFAP (reactive astrocyte marker), ID3, CD44 (Ast.3 marker). Complementary to **Fig. 3g**.

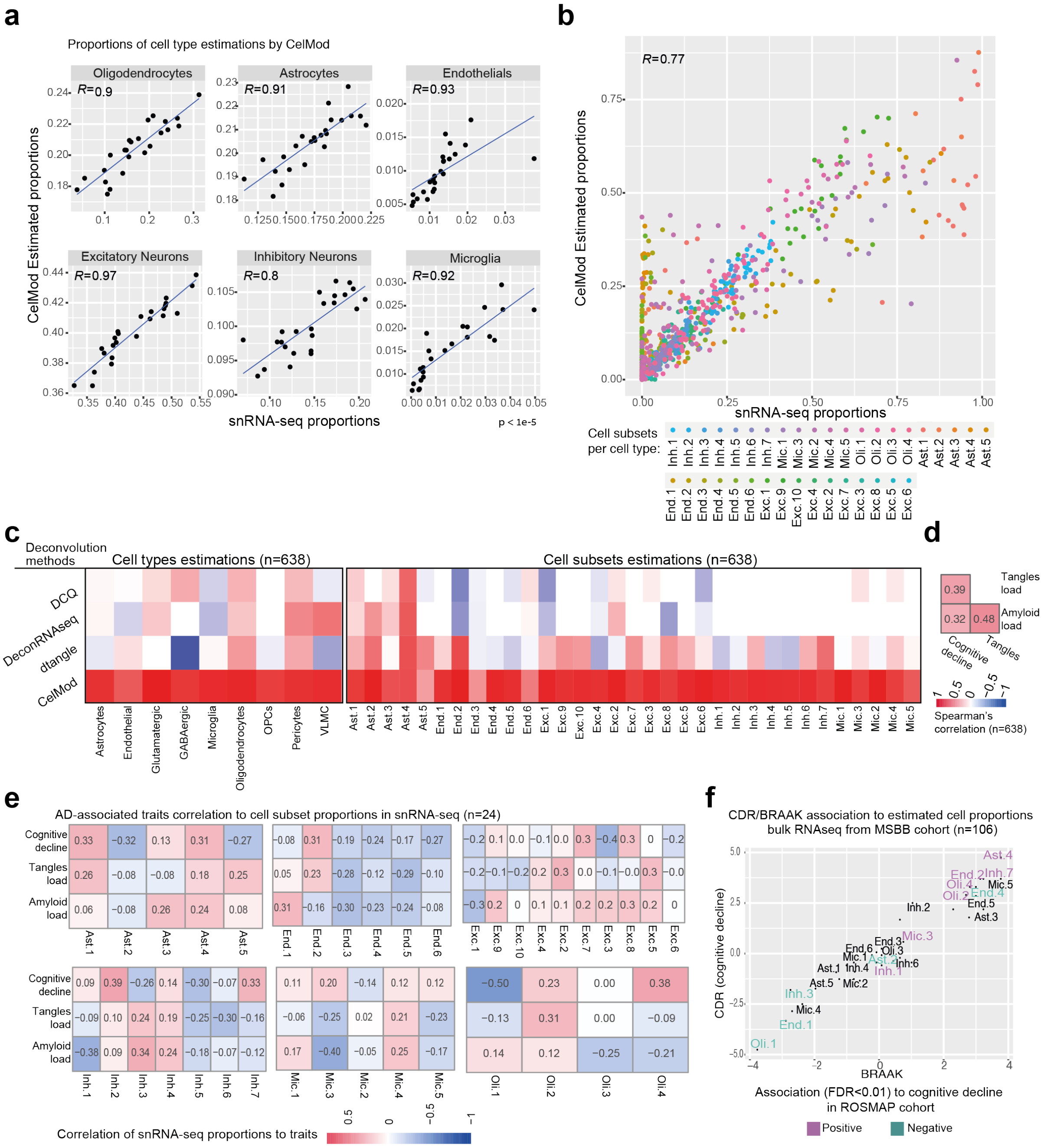


**Extended Data Figure 6. CelMod evaluation and comparison to other methods and datasets (Accompanying Fig. 4).** (**a,b**) Estimated cell type proportions by CelMod algorithm match snRNA-seq data. Scatter plots of the estimated proportions (Y-axis) compared to the snRNA-seq measured proportions (X-axis) across the 24 individuals, for all 8 major cell classes (in **a**), or for all cell subsets or topic model (average, colored by the cell subset, in **b**). (**c**) CelMod outperforms previous deconvolution-based methods in prediction accuracy. Spearman correlation scores (color scale) of the measured proportions in snRNA-seq of each cell type (left) and cell subsets (right), relative to the estimated proportions made by three previous models and CelMod. (**d**) Correlations between AD traits within our data. Pairwise correlations (color scale) of the 3 major AD-associated traits across 638 individuals. (**e**) Measured proportions of cellular subsets from snRNA-seq in 24 individuals correlate with AD pathology and cognitive decline. Correlation (color scale) of the proportions of each cell subset *within* each cell type (individual columns) to AD-traits (rows): β-amyloid burden, tau tangle pathology and cognitive decline. The proportions are calculated over the total number of nuclei per individual (n=24) within each cell type in the snRNA-seq data. (**f**) Comparison of trait associations to cellular proportions between the MSBB[32] and the ROSMAP[7] cohorts. Association scores of estimated proportions of cell subsets (proportions within each cell type) to two accepted measures of AD ranking: BRAAK stage which relates to the tangles load across the brain and the CDR which quantifies the level of cognitive decline. Association score = -log(FDR)*sign(beta), from multivariable linear regression analysis corrected for age, sex and RIN values. Cell subset names are colored by the significant associations to the cognitive decline rate in the 638 ROSMAP cohort.

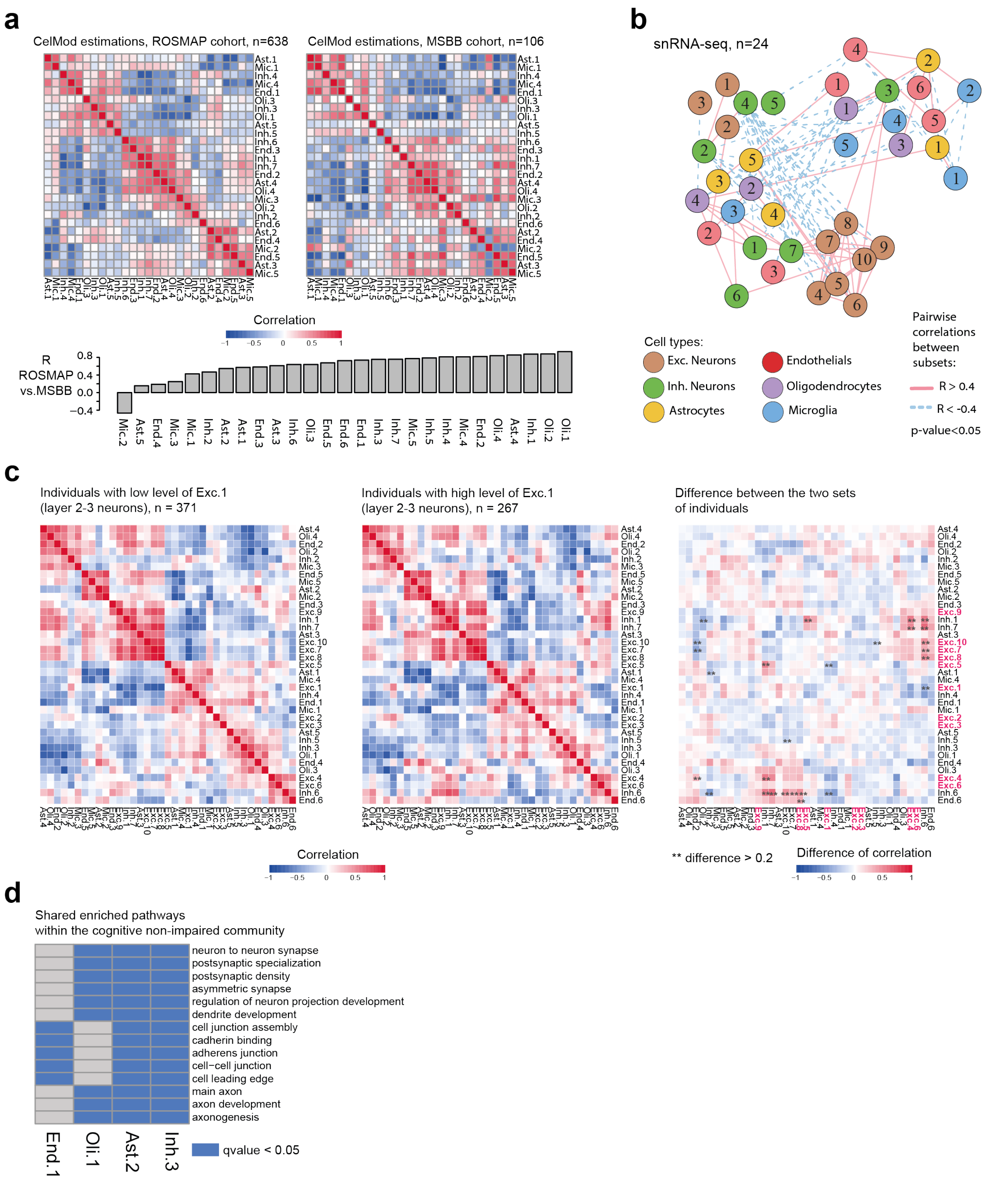


**Extended Data Figure 7. Evaluation of multi-cellular communities (Accompanying Fig. 5, 6 and 7).** (**a**) Similar structure of coordinated changes between proportions of cell states and subtypes across individuals found in two independent cohorts. Pairwise Spearman correlation coefficients of the proportions of cell states and subtypes across individuals estimated by CelMod in bulk RNA-seq data of ROSMAP[7] (left) and MSBB[32] (right) cohorts. Bottom: Correlation between the pairwise associations within the ROSMAP and the MSBB cohort per cell subset. (**b**) A network of cellular subsets reveals coordinated variation across individuals in multiple cell types. Network of coordinated and anti-coordinated cell subsets (nodes). Edges between pairs of subsets with significantly correlated proportions across individuals ($r>0.4$, p-value $< 0.05$, solid red line) or anti-correlated ($r<-0.4$, dashed blue line) based on snRNA-seq proportions (n=24). Celmod based network in **Fig. 6b**. Nodes are colored by the cell type and numbered by the subset as in **Fig. 2a and Fig 3. a,d,h**). (**c**) Coordinated changes in proportions of cell states and subtypes across individuals is independent of the cortical layer, except for excitatory neurons. Pairwise Spearman correlation coefficient of the CelMod proportions of all cell states and subtypes across individuals with low levels of Exc.1 (n=371 individuals, left) or high levels of Exc.1 (n=267 individuals, middle). Right: The differences in pairwise correlations between the Exc.1-high and Exc.1-low groups of individuals. Showing the partition mainly affects excitatory neurons (red). (**d**) Shared pathways within the cognitive non-impaired community. Enriched pathways (hypergeometric test, qvalue<0.05, blue) in up-regulated genes within each cell subset that are negatively associated with cognitive decline rate (End.1, Oli.1, Ast.2, and Inh.3). Displaying shared enriched pathways between at least three subsets within the cognitive *non-impaired* community.

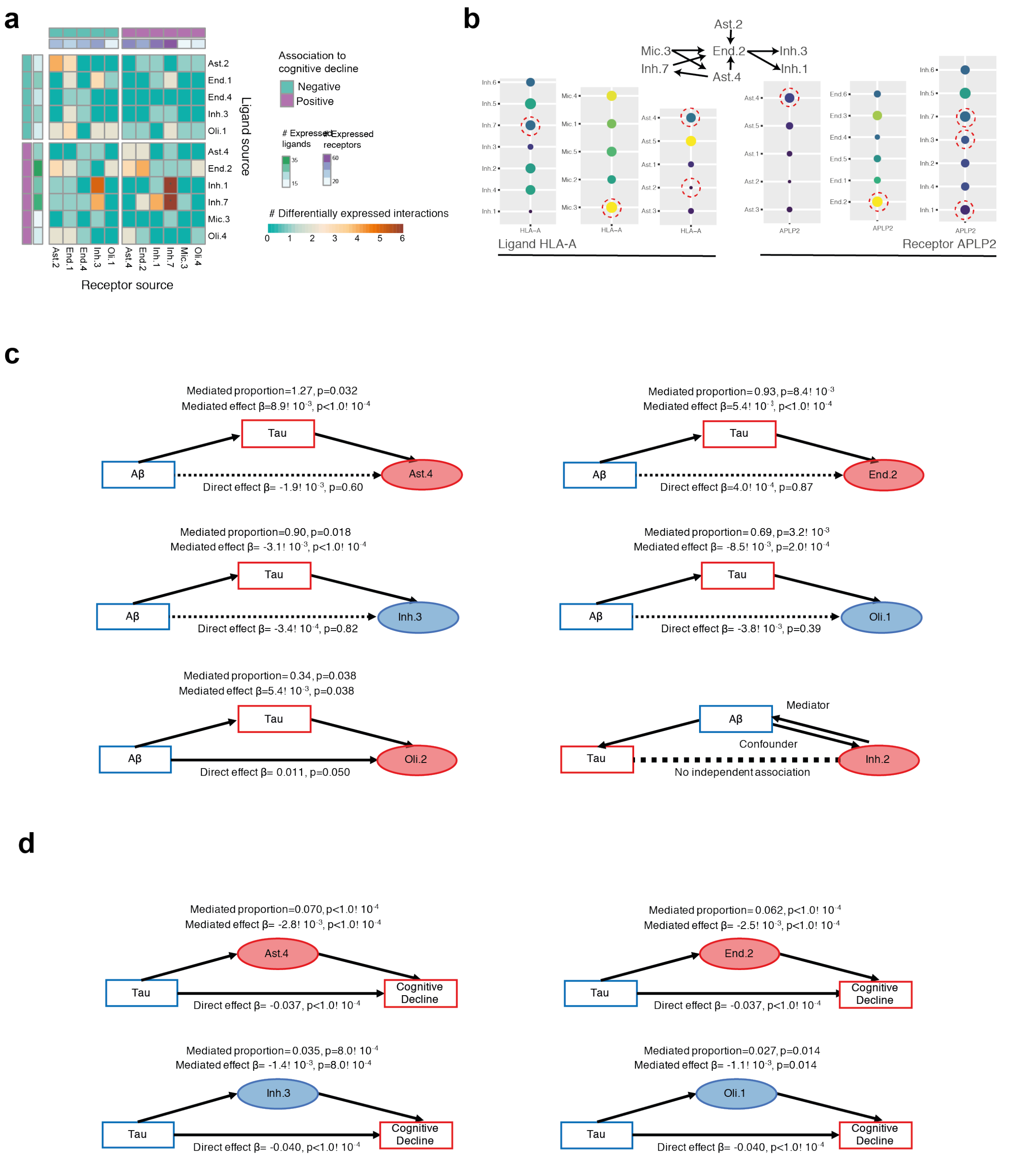


**Extended Data Figure 8. Signaling within and between multi-cellular communities and causal modeling (Accompanying Fig. 7) (a)** Cell subsets positively associated with cognitive decline have an increased expression of ligand-receptor pairs compared to the negatively associated subsets. For each pair of cell subsets showing the numbers (color scale) of ligand-receptor pairs (LRP, row: ligand, column: receptor) where both the ligand and the receptor are differentially expressed in the relevant subsets. Top and side bar marking: subsets positively associated with cognitive decline (purple) or negatively (turquoise), and the general number of ligands and receptors expressed (color scale). (**b**) Expression of the HLA-A - APLP2 ligand-receptor pair across subtypes of different cell types. Dot plot of the mean expression level in expressing cells (color) and percent of expressing cells (circle size) of HLA-A and of APLP2 across subsets of selected cell types. (**c**) Mediation analysis results showing tangle pathology burden (tau) is predicted to be upstream of changes in proportion of Inh.3, Oli.1, Oli.2, Ast.4, and End.2, but not Inh.2. (**d**) Mediation analysis results showing partial effect of changes in proportion of Inh.3, Olig.1, Ast.4, and End.2 on cognitive decline independent of tau pathology burden.
